# Liver Immunity-Guided Discovery of Capsule-Permeable Protein Antigens Against Bacterial Sepsis

**DOI:** 10.64898/2026.11.12.765860

**Authors:** Qian Jin, Yanrong Wu, Haiteng Deng, Wenwen Zeng, Jeremy S. Brown, Juanjuan Wang, Jing-Ren Zhang

**Author notes:** Corresponding author. (JW); (J-RZ). These authors contributed equally to this work (Qian Jin, Yanrong Wu).

## Abstract

The capsules of invasive bacteria shield underlying antigens, limit immune clearance and hinder vaccine noncapsular antigen discovery. To address this, we developed Liver-based Vaccine Antigen Capture (L-VAC), a functional platform integrating antibody-mediated screening with *in vivo* visualization of hepatic bacterial clearance in mice. Applying L-VAC to *Streptococcus pneumoniae* identified multiple capsule-permeable proteins that induce rapid hepatic clearance of blood-borne bacteria. Immunization with these proteins, especially in a multivalent formulation, conferred broad cross-serotype protection. Spatial analyses revealed that only the proteins localized at or near the capsular surface are targetable for vaccine-induced pathogen capture in the liver sinusoids. These findings establish L-VAC as a sensitive and reliable approach for identifying protective protein antigens capable of overcoming capsule barriers, offering a promising strategy to develop broadly protective vaccines against multidrug-resistant encapsulated pathogens.

**One Sentence Summary:** A liver-based antigen capture platform identifies capsule-permeable protein antigens that confer cross-serotype protection against bacterial sepsis.

## INTRODUCTION

Invasive bacterial infections are a major cause of human morbidity and mortality, yet effective vaccines are lacking for most of the top impactful pathogens, particularly for those “superbugs” with extreme antimicrobial resistance (*1, 2*). A central challenge in bacterial vaccine development is the identification of protective antigens, compounded by the lack of *in vivo* markers to quantify vaccine-induced pathogen elimination during infection. Genome-based approaches have enabled the identification of conserved extracellular proteins in many pathogenic bacteria (*3–6*), but protein-based vaccines are currently only available for serogroup B *Neisseria meningitidis* (*7, 8*). The efficacy of vaccines relies on two functional aspects: 1) vaccine-induced antibody and/or T-cell responses, and 2) vaccine-elicited pathogen elimination upon pathogen entry (*9*). While the first aspect has been developed well, the second—translating immune responses into effective pathogen clearance—remains a major bottleneck. Besides measuring survival of model animals, current vaccine evaluation systems mostly depend on *in vitro* assessments of vaccine-induced immune responses, including antibody titer, antibody-mediated neutralization, and opsonophagocytosis of pathogens (*10, 11*). However, these parameters do not always predict how vaccine-elicited immune responses translate into the host’s capacity for pathogen clearance during infection, as many vaccine candidates induce strong antibody responses but fail to confer anticipated protection in animal models.

Many invasive bacteria are coated by a protective layer of extracellular or capsular polysaccharides (CPSs), including the top five bacterial causes of human mortality (*2, 12*). CPSs are the protective antigens in the current vaccines for *Streptococcus pneumoniae, Haemophilus influenzae* and *N. meningitidis* (*13, 14*). Capsule-based vaccines confer effective immunity against the covered serotypes, but are unable to control infections of non-vaccine serotypes of the same pathogens. For instance, the decline in vaccine-covered serotypes of *S. pneumoniae* has accompanied the expansion of non-vaccine serotypes since the introduction of pneumococcal polysaccharide-conjugate vaccines, a phenomenon known as serotype replacement (*15, 16*). Recent surveillance studies indicate that non-vaccine serotypes now account for a substantial proportion of pediatric invasive pneumococcal disease (IPD) in many regions worldwide (*17, 18*). Additionally, complex manufacturing processes contribute to high production costs, limiting vaccine accessibility in resource-limited settings (*19*). Furthermore, protection against some vaccine-covered serotypes remains suboptimal. For example, serotype-3 pneumococci continue to cause pediatric IPD despite the inclusion of serotype-3 CPS in pneumococcal conjugate vaccines (*20–22*). Protein-based vaccines have been extensively explored as alternatives to capsule-based vaccines. Many pneumococcal proteins have been investigated as vaccine candidates (*23–26*). Because individual proteins typically induce relatively weaker protection, multicomponent protein formulations have also been explored (*27–32*). However, no protein-based vaccine has yet been licensed for pneumococcal disease prevention.

Our recent studies have revealed that the liver is the central immune organ responsible for antibody-mediated immunoprotection against pneumococcal septic infection. Vaccination with capsule-based vaccines engages the liver-resident macrophage Kupffer cells (KCs) and sinusoidal endothelial cells (LSECs) to rapidly capture and kill the antibody-opsonized bacteria in the bloodstream (*33*). Building on this mechanism, we previously proposed a liver-based framework to functionally evaluate the protective efficacy of vaccines against multiple encapsulated bacteria (*34*). In this study, we developed the liver-based vaccine antigen capture (L-VAC) approach to functionally screen protective protein antigens that enable hepatic capture of blood-borne encapsulated bacteria. Using L-VAC, we identified multiple pneumococcal protein antigens that induce hepatic clearance of blood-borne pneumococci across serotypes and confer cross-serotype protection.

## RESULTS

### Antibodies to non-capsular antigens promote the hepatic clearance of encapsulated pneumococci

While capsules shield the underlying non-capsular antigens from host immune recognition, vaccination with certain pneumococcal proteins or acapsular bacteria can nevertheless confer protection (*35, 36*). To investigate this paradox, we took advantage of a bacterial clearance mouse model established in our recent studies (*37, 38*). An unencapsulated mutant of serotype-4 *S. pneumoniae* TIGR4, a high-virulence strain resistant to hepatic capture in mice (*39*), was inactivated and used as a whole cell vaccine (WCV) to immunize mice (**Fig. 1A**). Following challenge with wild-type TIGR4, vaccinated mice rapidly cleared *S. pneumoniae* from the bloodstream resulting in a 50% clearance time (CT_50_) of 2.6 min, whereas naïve mice showed stable bacteremia over the first 30 min (**Fig. 1B**). After 30 min, only 13% of the inoculum was detected in vaccinated mice, whereas 85% of the inoculum bacteria remained detectable in naïve mice (**Fig. 1C**), demonstrating the effective vaccine-induced bacterial elimination. Consistently, all vaccinated mice survived infection with an otherwise lethal dose (**Fig. 1D**). WCV-immunized mice also showed little to no detectable bacteremia from 12 h (**Fig. 1E**). In contrast, naïve mice developed severe bacteremia, with only a transient decline of bacteremia due to the splenic clearance (*39*). Similar vaccine-induced protection was also observed in BALB/c mice (**Fig. S1A-D**). Together, these data demonstrate that vaccination with non-capsular antigens induced robust protection against encapsulated pneumococci.

**Fig. 1.**
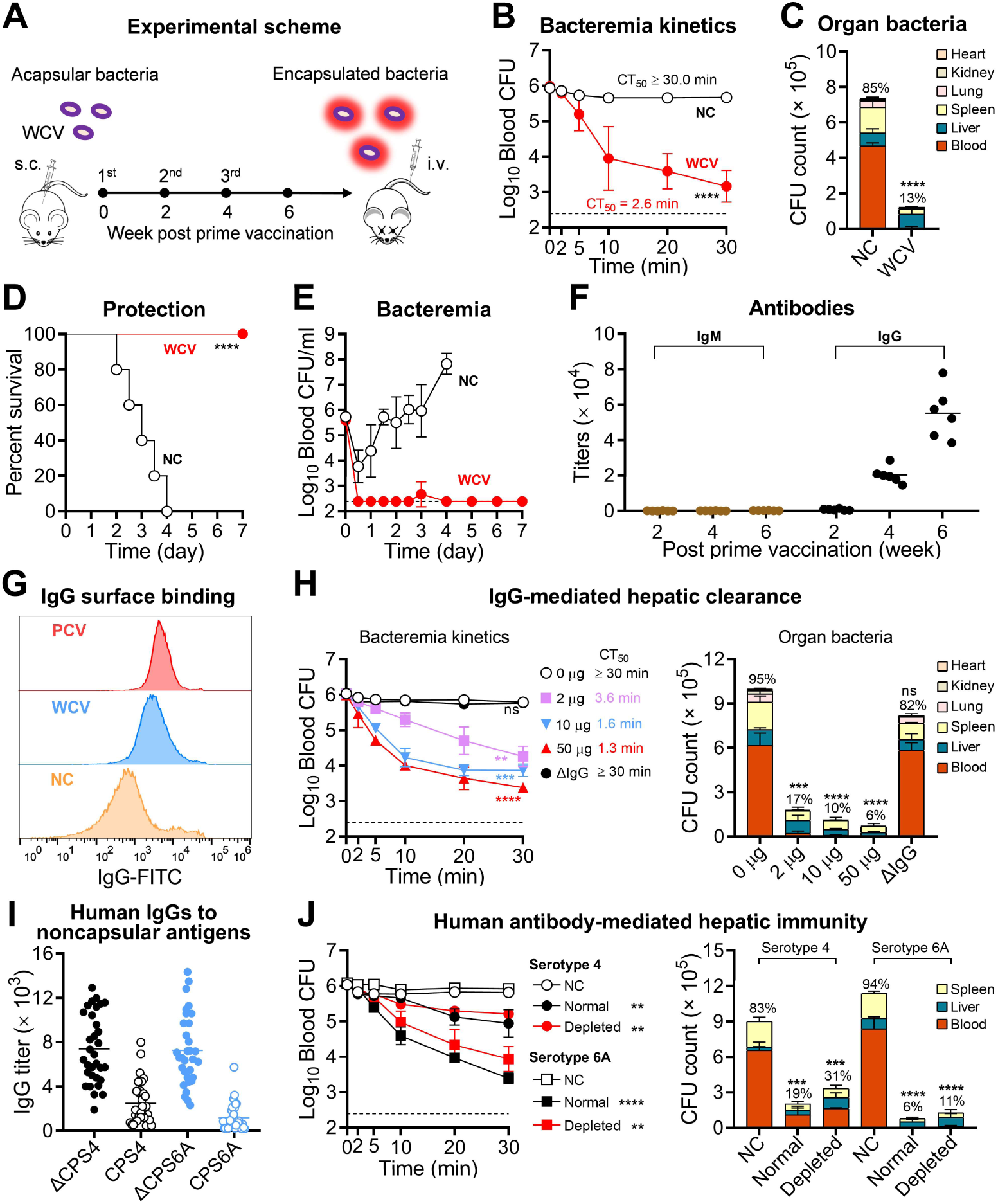
The immunoprotection of non-capsular antigens against encapsulated pneumococci. (**A**) Experimental scheme for vaccination and challenge. CD1 mice were subcutaneously immunized three times with the whole-cell vaccine (WCV) of acapsular TIGR4, and intravenously challenged with 10^6^ CFU of encapsulated TIGR4 two weeks after the final boost. **(B** and **C)** The impact of vaccination on the bacterial clearance from the bloodstream in the liver was assessed by enumerating CFUs of blood (B) and major organs (C) of vaccinated (WCV) and control (NC) mice post i.v. infection with 10^6^ CFU of TIGR4. Dotted line indicates the detection limit. CT_50_, 50% clearance time; CT_50_ was designated as 30 min if less than half of the inoculum was cleared from the blood at 30 min. The percentage of viable bacteria remaining at 30 min relative to the initial inoculum is shown above the bar. n = 3-6. **(D** and **E)** Vaccine-elicited protection is presented as the survival rate (D) or bacteremia kinetics (E) of mice after i.v. infection with 10^6^ CFU of TIGR4. n = 10. **(F)** Serum IgG and IgM titers were measured by ELISA at 2, 4, and 6 weeks post prime immunization. n = 6. **(G)** IgG binding to the encapsulated TIGR4 was detected by flow cytometry. **(H)** IgG-mediated clearance of encapsulated TIGR4 was determined by enumerating CFU in the blood (left panel) and major organs (right panel) post i.v. infection with 10^6^ CFU of pneumococci that were pre-incubated with various amounts of purified IgG or IgG-depleted serum (ΔIgG). n = 3. **(I)** IgG titers against acapsular pneumococci (ΔCPS4 and ΔCPS6A) or purified CPS4 and CPS6A were measured in normal human sera by ELISA. n = 32. **(J)** Hepatic clearance of pneumococci promoted by normal human sera or human sera depleted of CPS-specific IgG in FcγRs-humanized mice. Bacteremia kinetics (left panel) and organ bacteria at 30 min (right panel) were detected post i.v. infection with 10^6^ CFU of serotype-4 or serotype-6A pneumococci pre-opsonized with 10 μl of human sera with or without depletion of CPS-specific antibodies. NC, no serum. n = 3. The representative data of at least two independent experiments are presented as mean ± SD, with statistical differences determined by two-way ANOVA with Tukey’s multiple comparisons test (B, left panel in H and J), unpaired *t* tests (C) and multiple *t* tests (right panel in H and J), and log-rank (Mantel-Cox) test (D). ns, no significant; \*\**P* < 0.01; \*\*\**P* < 0.001; \*\*\*\**P* < 0.0001.

Since antibodies are essential for polysaccharide vaccine-induced hepatic trapping of encapsulated bacteria (*33*), we assessed their role in WCV-induced immunity. WCV immunization induced high levels of IgG antibodies, dominated by the IgG1 and IgG2b subclasses, with negligible IgM (**Fig. 1F**; **Fig. S1E-F**). Flow cytometry showed robust IgG binding to encapsulated pneumococci (**Fig. 1G**). We next purified IgG from WCV immune serum to directly assess its contribution to bacterial clearance (**Fig. S1G**). Pre-opsonization of TIGR4 with purified IgG significantly accelerated bacterial clearance from the blood circulation of naïve mice in a dose-dependent manner (**Fig. 1H**, left panel), and resulted in substantial bacterial killing (**Fig. 1H**, right panel). These results demonstrate that IgG antibodies elicited by non-capsular antigens are able to recognize encapsulated pneumococci and promote their rapid hepatic clearance.

It is well-known that natural exposure to *S*. *pneumoniae* through colonization or infection induces antibodies against capsular and non-capsular antigens in humans (*40*). We thus tested whether human antibodies to non-capsular antigens could promote pneumococcal clearance in FcγRs-humanized mice (*33*). Antibodies to encapsulated and acapsular pneumococci were detected in normal human sera (**Fig. 1I**). After depletion of CPS-specific antibodies (**Fig. S1H**), human sera retained significant activity in promoting hepatic clearance of serotype-4 and -6A pneumococci in FcγRs-humanized mice (**Fig. 1J**), consistent with previous evidence that pooled human IgG depleted of anti-capsular antibodies can protect against *S. pneumoniae* sepsis (*41*). These findings demonstrate that human antibodies to non-capsular antigens can also play an important role in the clearance of encapsulated pneumococci.

Finally, we assessed whether non-capsular antigens from other encapsulated bacteria could elicit similar protective immunity by immunizing mice with inactivated capsule/O antigen-deficient K2 *Klebsiella pneumoniae*, a major antimicrobial-resistant pathogen (*2*) (**Fig. S1I**). Pre-opsonization of K1, K16, K20 and K47 *K. pneumoniae* strains with the resulting immune serum significantly accelerated their hepatic clearance (**Fig. S1J-K**), indicating that antibodies to non-capsular antigens can promote the clearance of encapsulated bacteria beyond *S. pneumoniae*.

### Liver-based vaccine antigen capture (L-VAC) identified *in vivo* protective capsule-permeable protein (CPP) antigens

The robust hepatic clearance observed in WCV-immunized mice strongly suggested that antibodies binding to certain non-capsular antigens or capsule-permeable proteins (CPPs) can promote hepatic capture of encapsulated bacteria. To identify these CPPs, we developed liver-based vaccine antigen capture (L-VAC), an antibody-guided surface protein discovery approach, using *S. pneumoniae* as a model (**Fig. 2A**). Encapsulated pneumococci were incubated with acapsular WCV immune serum to enrich IgG antibodies bound to CPPs on intact bacteria (**Fig. 2B**); the resulting antibody-antigen complexes were then pulled down with Protein A/G beads. After exclusion of ribosomal proteins, mass spectrometry identified 90 proteins enriched at least 5-fold in WCV immune-serum group relative to the control-serum group (**Table S1**). Notably, the top 20 hits were each enriched at least 20-fold, and all were proteins known or predicted to associate with the cell wall (CbpA, Eno and PspA), pili (RrgA, RrgB and RrgC) or cell membrane (AmiA, GlpF, LivJ, MltG, PBP1a, PiaA, PnrA, PrsA, PsaA, SP0148, SP0282, SP1241, SP1926 and SP2101) (**Table S2**). The preferential enrichment of cell-envelope-associated proteins suggested that L-VAC captures protein antigens that are antibody-accessible on intact encapsulated bacteria.

**Fig. 2.**
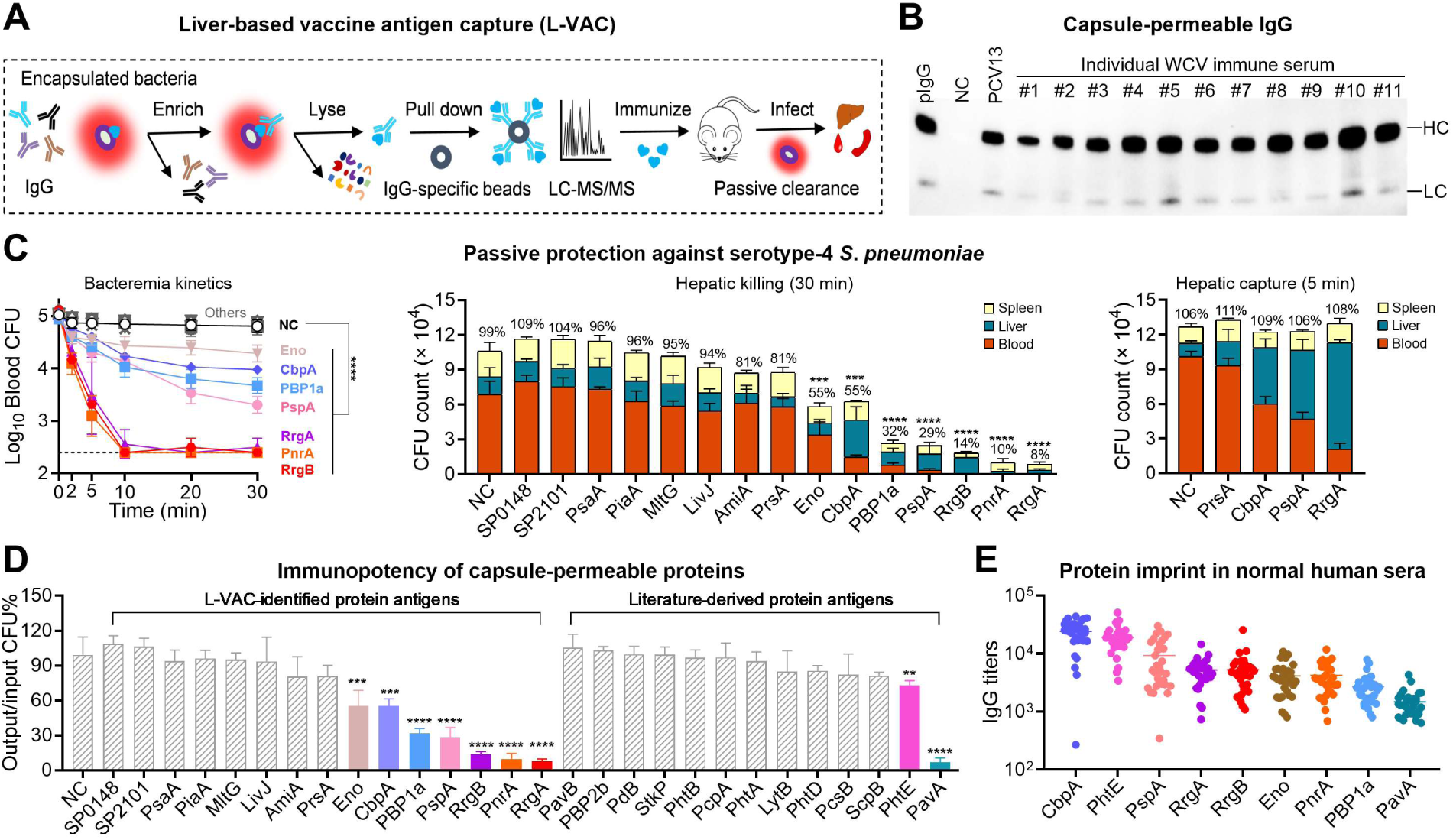
Identification of capsule-permeable proteins by the liver-based vaccine antigen capture (L-VAC). **(A)** Schematic illustration of L-VAC to identify pneumococcal CPPs using WCV immune serum. **(B)** Detection of bacterium-bound IgG by immunoblotting with a rabbit anti-mouse IgG secondary antibody. TIGR4 pneumococci (10^5^ CFU) were incubated with immune serum from mice immunized with WCV, PCV13 or adjuvant control (NC), followed by SDS-PAGE and immunoblotting. Purified IgG (pIgG) from WCV immune serum was used as a positive control. HC, IgG heavy chain; LC, IgG light chain. **(C)** Functional validation of the top hits using an immune serum-mediated passive clearance assay. Immune sera were collected from mice after three immunizations with recombinant proteins; TIGR4 pneumococci (10^5^ CFU) were opsonized with 20% immune sera and i.v. injected into naïve mice to assess bacterial clearance from the bloodstream (left panel), bacterial killing (center panel) and hepatic capture (right panel). NC, adjuvant control. n = 3-6. **(D)** Protein antigen-induced bacterial killing in mice is presented as the percent ratio between the total viable bacteria (in the blood, liver and spleen) at 30 min post infection and the inoculum (10^5^ CFU). The left and right panels show the protein hits of this work and the previously reported pneumococcal proteins, respectively. NC, adjuvant control. n = 3-5. **(E)** IgG titers against nine CPPs in normal human sera. n = 32. The representative data of at least two independent experiments are presented as mean ± SD, with statistical differences determined by two-way ANOVA with Tukey’s multiple comparisons test (left panel in C), and multiple *t* tests (center and right panels in C). \*\*\**P* < 0.001; \*\*\*\**P* < 0.0001.

We next evaluated the ability of the top L-VAC hits to induce hepatic bacterial clearance. Of the 20 most highly enriched candidates, five (GlpF, RrgC, SP0282, SP1241 and SP1926) failed to yield soluble recombinant proteins, likely due to their predicted transmembrane segments. We therefore evaluated the remaining 15 candidates by immunizing mice with their recombinant proteins. In the liver-based bacterial clearance model, immune sera against 7 of the 15 proteins (46.7%)—CbpA, Eno, PBP1a, PnrA, PspA, RrgA and RrgB—significantly promoted hepatic clearance of encapsulated pneumococci, though the extent of protection varied between the antigens (**Fig. 2C**). Immune sera against PnrA, RrgA and RrgB showed the strongest activity, reducing bloodstream CFU to near or below the detection limit by 10 min after i.v. inoculation with 10^5^ CFU of serum-opsonized pneumococci (**Fig. 2C**, left panel), while immune sera against CbpA, Eno, PBP1a, and PspA produced weaker but still significant clearance activity. Enumeration of total viable bacteria recovered from mice showed a marked reduction in bacteria pre-opsonized with anti-PnrA, -RrgA, and -RrgB serum, with less than 20% of the inoculum viable at 30 min (**Fig. 2C**, center panel). To a lesser extent, pre-opsonization with the immune sera against CbpA, Eno, PBP1a, and PspA also led to significant reductions in bacterial burden. Consistent with the dominant role of the liver in capturing anti-capsule antibody-opsonized bacteria (*33*), bacteria cleared from the bloodstream were detected predominantly in the liver at 5 min post-infection (**Fig. 2C**, right panel). These functional validation experiments demonstrate that L-VAC can identify CPPs capable of promoting the *in vivo* clearance of encapsulated pathogens.

We also assessed the hepatic clearance activity induced by 13 additional previously reported protective antigens of *S. pneumoniae*, including LytB (*25*), PavA (*6*), PavB (*42*), PBP2b (*31*), PcpA (*43*), PcsB (*24, 31*), PhtA (*44*), PhtB (*44*), PhtD (*43, 44*), PhtE (*44*), pneumolysin (*45*), ScpB (*31*), and StkP (*24*). While opsonization with anti-PavA serum resulted in sterilizing clearance of TIGR4 pneumococci, anti-PhtE serum showed a much weaker but significant clearance activity (**Fig. 2D**; **Fig. S2A-E**). Under these conditions, pre-treatment with the immune sera against the remaining 11 proteins did not yield obvious hepatic clearance of TIGR4. Lastly, consistent with previous studies (*5, 46*), ELISA using human sera detected substantial levels of IgG antibodies recognizing all of the 9 CPPs (**Fig. 2E**), with the highest IgG titers found for CbpA, PhtE and PspA. These findings suggest that these CPPs are recognized by naturally acquired human antibodies.

### CPPs elicit cross-serotype protection against pneumococcal septic infections

We further assessed the cross-serotype immunity of the nine CPPs. Opsonization with the CPP-specific immune sera revealed marked differences in both bloodstream clearance and bacterial killing across serotypes 1, 3, 5, 6A, 8 and 15A. While immune sera against CbpA, Eno, PBP1a, PhtE, PspA and RrgA displayed cross-serotype clearance activity, the immune sera against RrgB, PavA and PnrA showed little or no hepatic clearance activity against these six serotypes tested (**Fig. 3A-B**; **Fig. S3**).

**Fig. 3.**
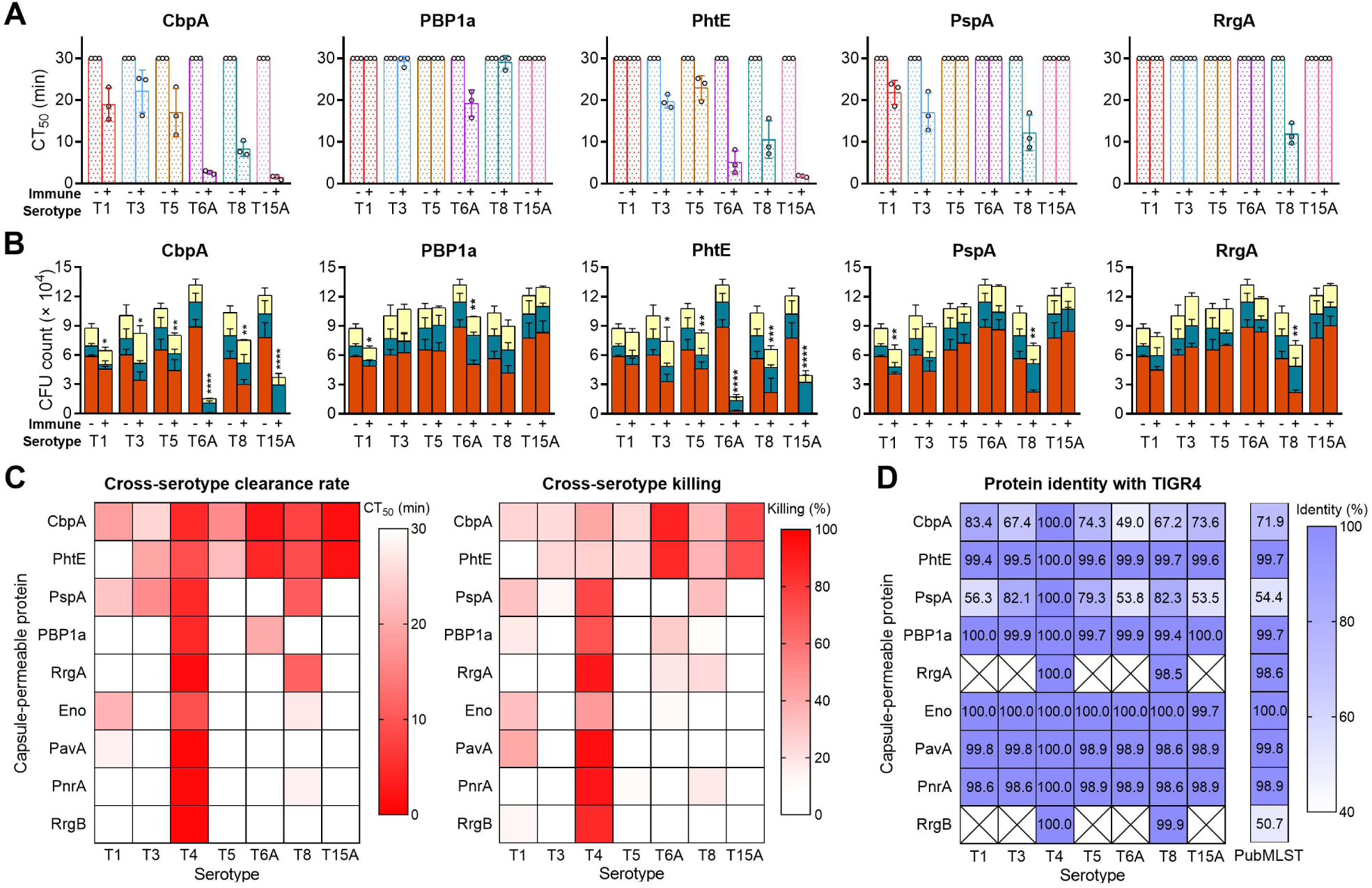
Cross-serotype pathogen clearance elicited by the capsule-permeable proteins. **(A)** CPP-induced clearance of serotypes 1, 3, 5, 6A, 8 and 15A pneumococci was assessed in naïve mice using immune serum-opsonized bacteria (10^5^ CFU) as in Fig. 2C. The data are presented as a CT_50_ value for each serotype as in Fig. 1B. CT_50_ was designated as 30 min if less than half of the inoculum was cleared from the blood at 30 min. -, adjuvant control; +, immune serum. n = 3. **(B)** The killing rate of immune serum-opsonized pneumococci in naïve mice is presented as viable bacteria (CFU count) in the blood (red), liver (blue) and spleen (yellow) at 30 min post i.v. inoculation. -, adjuvant control; +, immune serum. n = 3. **(C)** Cross-serotype immune potencies of the CPPs in inducing pathogen clearance from the blood (left panel) and killing (right panel) shown in (A) and (B) are summarized as heatmaps. n = 3-5. (D) Amino acid sequence identities of nine CPPs from the tested serotype 1, 3, 5, 6A, 8 and 15A strains relative to TIGR4, together with the median identities of 34,099 pneumococcal genomes in PubMLST, are summarized as heatmaps. The representative data of at least two independent experiments are presented as mean ± SD, with statistical differences determined by multiple *t* tests (B). ns, no significant difference; \**P* < 0.05; \*\**P* < 0.01; \*\*\**P* < 0.001; \*\*\*\**P* < 0.0001.

Among the nine CPPs, CbpA, PhtE and PspA showed the broadest immune coverage of tested pneumococcal serotypes (**Fig. 3A**; **Table S3**). As quantified by the 50% clearance time (CT_50_) of opsonized bacteria in the bloodstream, CbpA serum significantly accelerated the clearance of serotype-1, -3, -4, -5, -6A, -8 and -15A pneumococci from the bloodstream, despite substantial sequence variations in CbpA among strains (**Fig. 3D**; **Table S3**) (*47*). This broad activity is consistent with previous studies demonstrating the vaccine potential of CbpA, also known as PspC (*48–51*). PhtE showed the second broadest coverage, significantly accelerating clearance of all serotypes tested except serotype 1, and was highly conserved among these strains (**Table S3**). This finding is consistent with previous reports that PhtE, a pneumococcal surface-exposed histidine triad protein, induces cross-serotype immunity in mice (*44, 52, 53*). PspA immune serum significantly promoted clearance of serotypes 1, 3, 4 and 8, but not serotypes 5, 6A or 15A, consistent with its previously reported cross-serotype protective activity despite extensive sequence variation among strains (*54, 55*). However, only PspA from the serotype-3 and 4 strains belonged to Family 2 based on Clade Determination Region (CDR) identity (*56*) (**Table S3**), suggesting that the observed cross-serotype activity may be attributed to regions outside the CDR within the N-terminal alpha-helical domain (*54, 57*).

The other CPPs exhibited more restricted patterns of cross-serotype activity. PBP1a immune serum only promoted the clearance of serotype 6A in addition to serotype 4. Consistent with the strain-restricted distribution of the pilus locus (*58*), RrgA and RrgB showed limited but distinct cross-serotype activities. RrgA immune serum promoted clearance of serotypes 4 and 8, whereas RrgB activity was restricted to serotype 4 TIGR4 (**Fig. 3B**; **Fig. S3**). This difference may reflect their distinct structural positions and antibody accessibility within the pilus, with RrgA serving as an ancillary adhesin and RrgB forming the major pilus backbone. Although highly conserved across pneumococcal strains, Eno immune serum promoted significant clearance only of serotype 1 and serotype 4, indicating that sequence conservation alone does not predict functional coverage. Eno was not prioritized for subsequent vaccine development because of its potential human cross-reactivity (*59*). Despite their strong immune activities against TIGR4 (**Fig. 2C**), IgG to PavA and PnrA exhibited little or no clearance activity against the additional serotypes tested (**Fig. 3C**; **Fig. S3**). CPP-mediated bloodstream clearance was generally accompanied by increased bacterial killing. At 30 min after i.v. inoculation, CPP immune sera significantly reduced viable bacterial burdens, with killing patterns largely consistent with their clearance activity.

We next examined the population-level conservation of the nine CPPs using 34,099 pneumococcal genomes in PubMLST. Eno, PavA, PBP1a, PhtE, and PnrA were broadly conserved, with ≥90% query coverage in 93.3-99.8% of genomes and median amino acid identities of 98.9-100.0% (**Fig. 3D**; **Table S3**). In contrast, the TIGR4 CbpA and PspA sequences met these matching criteria in 52.6% and 28.7% of genomes, with median identities of 71.9% and 54.4%, respectively. RrgA and RrgB showed restricted distributions, matching approximately 24.3% of genomes, with median identities of 98.6% and 50.7%, respectively (**Fig. 3D**; **Table S3**). Thus, Eno, PavA, PBP1a, PhtE, and PnrA proteins are broadly conserved, whereas CbpA/PspA are more divergent and RrgA/RrgB show restricted matching. Notably, sequence conservation did not directly predict functional breadth, as the highly conserved Eno, PavA, and PnrA showed limited cross-serotype activity, whereas the more variable CbpA exhibited broad functional coverage. These complementary activity profiles suggested that combining selected CPPs could broaden serotype coverage.

### The combination of multiple CPPs enhances cross-serotype immunity

To improve protective efficacy, we combined five CPPs (CbpA, PBP1a, PhtE, PspA and RrgA) to generate a pentavalent protein vaccine (PentaV). These proteins were selected based on the complementary serotype coverage. While PentaV induced comparable levels of antigen-specific IgG to those of individual protein immunizations (**Fig. S4A**), opsonization with its immune serum yielded significantly higher levels of hepatic clearance of the PCV13 serotypes 4 and 6A, and non-PCV13 serotype 15A, as compared with the immune sera against individual proteins (**Fig. 4A**). Furthermore, PentaV showed a broader serotype coverage. PentaV-immunized mice efficiently cleared both PCV13-covered (1, 3, 4, 5, and 6A) and non-PCV13 serotypes (8 and 15A), although the magnitude of clearance varied among serotypes (**Fig. 4B**; **Fig. S4C)**. Taken together, these data show that the multivalent CPP vaccine was superior to the individual CPPs in promoting hepatic pathogen clearance and broadening serotype coverage.

**Fig. 4.**
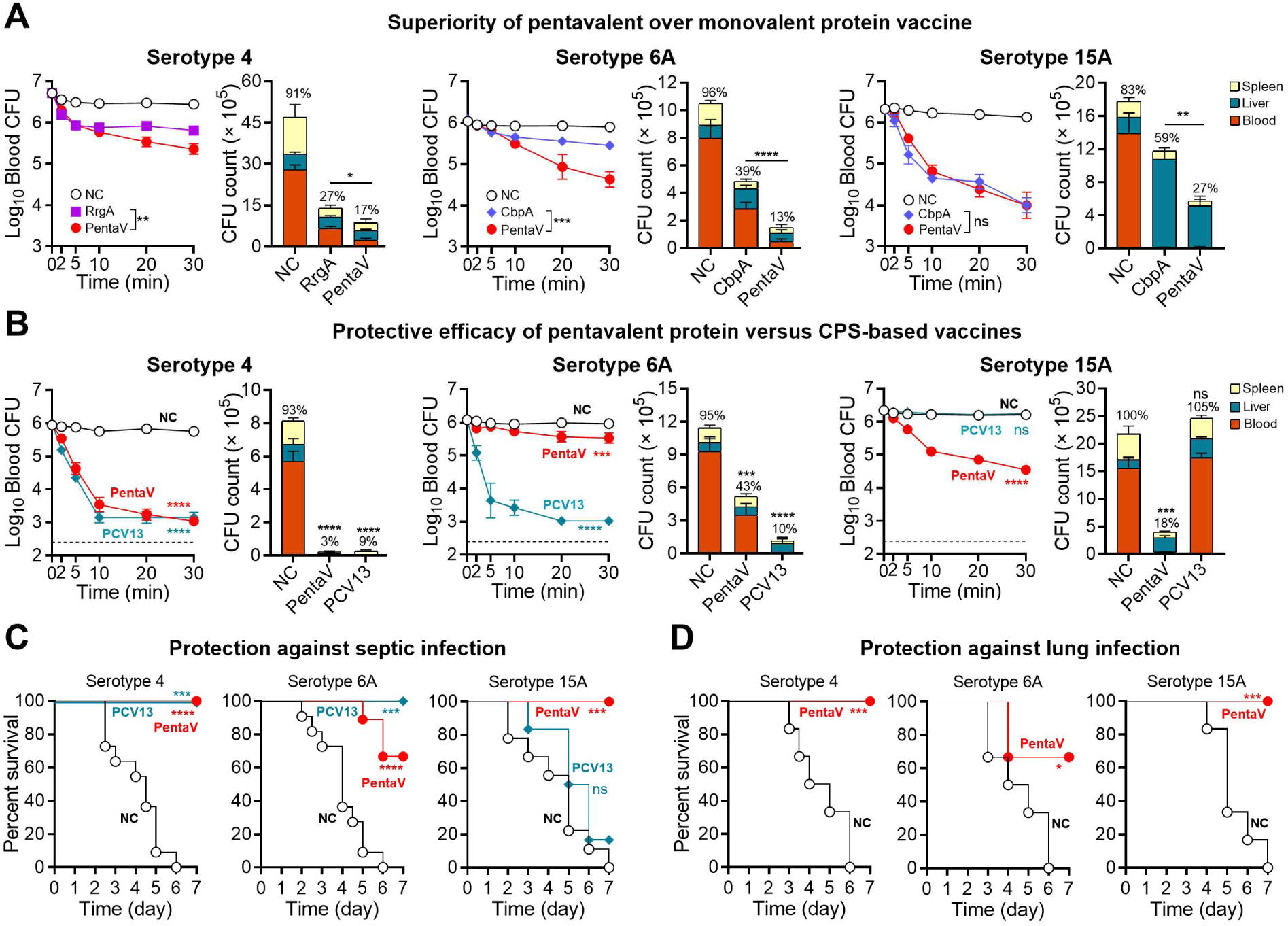
Cross-serotype protection of a pentavalent protein vaccine (PentaV). **(A)** The PentaV immune serum-mediated passive clearance of serotype 4 (5 × 10^6^ CFU), serotype 6A (10^6^ CFU), or serotype 15A (2 × 10^6^ CFU) pneumococci was assessed as in Fig. 2C, and presented as bacteremia dynamics (left panel) in the first 30 min and the viable bacteria in the blood, liver and spleen at 30 min post i.v. inoculation of serum-opsonized bacteria (right panel). CbpA and adjuvant sera were used as positive and negative controls, respectively. n = 3-6. **(B)** Cross-serotype bacterial clearance induced by active immunization with PentaV or PCV13. Vaccinated mice were i.v. infected with encapsulated pneumococci of serotypes 4 (10^6^ CFU), 6A (10^6^ CFU) or 15A (2 × 10^6^ CFU) to determine bacteremia kinetics (right panel) and organ bacteria at 30 min (left panel). NC, adjuvant control. n = 3. **(C)** The survival rate of PentaV-or PCV13-immunized mice in 7 days after i.v. infection with serotype-4 (1 × 10^6^ CFU), -6A (10^6^ CFU) or -15A (2 × 10^6^ CFU) pneumococci. NC, adjuvant control. n = 6-11. **(D)** The survival rate of PentaV-or PCV13-immunized mice in 7 days after intranasal (i.n.) infection with serotype-4 (2 × 10^6^ CFU), -6A (10^6^ CFU) or -15A (2 × 10^7^ CFU) pneumococci. NC, adjuvant control. n = 6. The representative data of at least two independent experiments are presented as mean ± SD, with statistical differences determined by two-way ANOVA with Tukey’s multiple comparisons test (left panel in A and B) and multiple *t* tests (right panel in A and B) and log-rank (Mantel-Cox) test (C, D). ns, no significant difference; \**P* < 0.05; \*\**P* < 0.01; \*\*\**P* < 0.001; \*\*\*\**P* < 0.0001.

We then compared the vaccine efficacy of PentaV to PCV13 using infection models with two PCV13 serotypes (4 and 6A) and one non-PCV13 serotype (15A). In both PentaV-and PCV13-immunized mice, more than 90% of serotype-4 pneumococci were cleared from the bloodstream by 30 min post-infection (**Fig. 4B**; **Fig. S4B**). Vaccination with PentaV induced a relatively lower level of hepatic clearance of serotype 6A compared to vaccination with PCV13, but showed significant efficacy against the non-PCV13 serotype 15A, whereas PCV13, as expected, had no impact. Consistent with the blood clearance data, immunization with PentaV and PCV13 fully protected mice from lethal serotype 4 sepsis, but PentaV induced significant but partial protection against serotype 6A although all PCV13-immunized mice survived (**Fig. 4C**; **Fig. S4D**). Immunization with PentaV but not PCV13 resulted in complete survival of serotype 15A-infected mice. Vaccination with PentaV also provided cross-protective immunity in murine lethal lung infection models with serotypes 4 and 15A (complete protection) or 6A (partial protection) (**Fig. 4D**; **Fig. S4E**). These data indicate that a multivalent CPP vaccine can induce effective and broad immunity against invasive pneumococcal diseases.

### CPP immunization promotes pathogen capture by liver macrophages and endothelial cells

To determine the mechanisms of the CPP-induced pathogen capture, we used intravital microscopy (IVM) to characterize the dynamic behavior of pneumococci in the liver sinusoids of PentaV-immunized mice. While pneumococci passed freely through sinusoidal vessels of naïve mice, PentaV-immunized mice rapidly immobilized circulating pneumococci to the liver sinusoidal vessel walls (**Fig. 5A**, PentaV; **Movie S1**; **Fig. S5A**). The immobilized bacteria were predominantly associated with the liver-resident macrophages, Kupffer cells (KCs), and to a lesser extent, with the liver sinusoidal endothelial cells (ECs). A similar pattern of hepatic capture was observed during active clearance in WCV-immunized mice and passive clearance in FcγRs-humanized mice (**Fig. 5A**, WCV; **Movie S1**; **Fig. S5B**). This result showed that KCs and ECs co-execute the CPP-induced pathogen capture in the liver of mice.

**Fig. 5.**
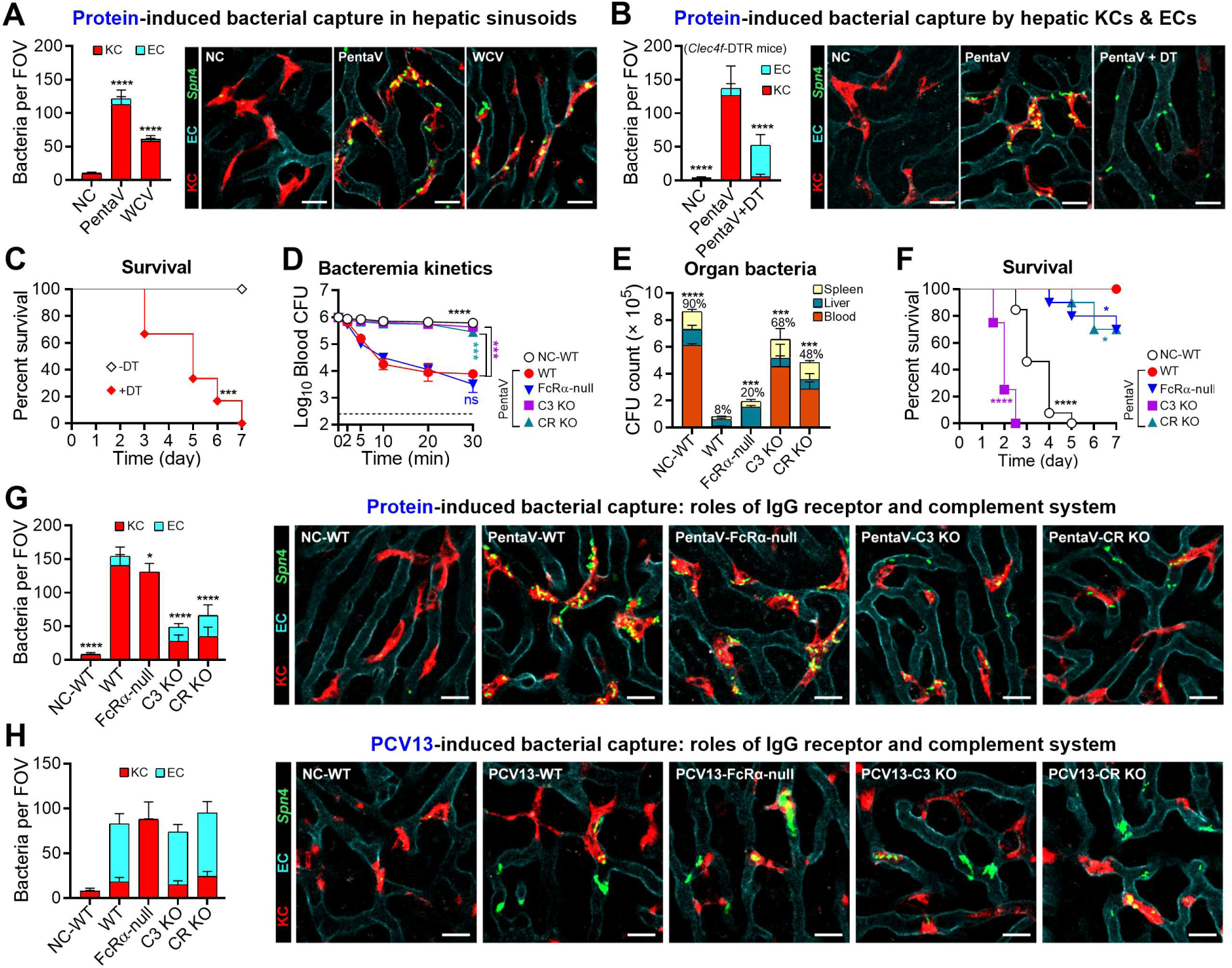
Protein antigen-induced pathogen capture by hepatic sinusoidal macrophages and endothelial cells. **(A)** Protein antigen-activated pneumococcal capture in the liver vasculatures. Mice immunized with PentaV were infected with 5 × 10^7^ CFU of serotype-4 pneumococci (green) and visualized for bacterial capture in the livers by IVM. Representative images of KC- and EC-associated bacteria (red) and ECs (cyan) were quantified by analyzing 6-12 random fields of view (FOV) (left panel). NC, adjuvant control; KC, Kupffer cell; EC, endothelial cell. Scale bar, 20 μm. n = 6-12 FOVs. Additional data are available in **Movie S1**. **(B** to **C**) Requirement for KCs in protein immune serum-mediated bacterial clearance. Serotype-4 pneumococci pre-opsonized with PentaV immune serum were intravenously injected into *Clec4f*-DTR mice treated with (+DT) or without (−DT) diphtheria toxin for IVM visualization (B; 5 × 10^7^ CFU, 100% serum) as in (A), and animal survival (C; 3 × 10^6^ CFU, 50% serum). Additional data are available in **Movie S2**. Scale bar, 20 μm. n = 6-12 in (B) and 6 in (C). **(D** to **G)** The roles of Fcγ receptors and the complement system in protein-induced bacterial clearance. Serotype-4 pneumococci pre-opsonized with PentaV immune serum were i.v. injected into mice lacking all the Fcγ receptors (FcRα-null), complement C3 (C3 KO) or C3 receptors (CR KO) for bacterial early clearance (D and E; 1 × 10^6^ CFU, 20% serum), animal survival (F; 1 × 10^6^ CFU, 50% serum) and IVM visualization (G; 5 × 10^7^ CFU, 100% serum) as in (A). Additional data are available in **Movies S3** and **S4**. n = 3 in (D and E), 8-12 in (F) and 6-12 in (G). **(H)** The roles of Fcγ receptors and the complement system in PCV13-induced bacterial capture. IVM was performed as in (G) except for using PCV13 immune serum for opsonization. n = 6-12. The representative data of at least two independent experiments are presented as mean ± SD, with statistical differences determined by two-way ANOVA with Tukey’s multiple comparisons test (D), multiple *t* tests (A, B, E and G) and log-rank (Mantel-Cox) test (C and F). ns, no significant difference; \**P* < 0.05; \*\*\**P* < 0.001; \*\*\*\**P* < 0.0001.

We further assessed the relative contributions of KCs and ECs to CPP-induced pathogen capture in the liver sinusoids of mice by IVM. IVM revealed a 61% reduction in vascular immobilization of PentaV serum-opsonized pneumococci after KC depletion (**Fig. 5B**, PentaV; **Movie S2**). Interestingly, the absence of KCs significantly enhanced bacterial capture by ECs. KC-deficient mice still showed significant bacterial capture as compared to WT mice, and all the immobilized bacteria were tethered to ECs (**Fig. 5B**, PentaV+DT; **Movie S2**; **Fig. S5C**). These findings indicate that KCs outcompete ECs in realizing CPP-induced pathogen capture. Consistently, KC-deficient mice were more susceptible to pneumococcal septic infection (**Fig. 5C**).

To define the molecular mechanisms of the CPP-induced pathogen capture, we tested the impact of Fcγ receptor deficiency on the phenotype. Mice lacking all four FcγRs (FcRα-null) displayed a modest but significant impairment in bacterial clearance and host survival (**Fig. 5D-F**; **Fig. S5D-E**). These mice showed no EC-immobilized bacteria (**Fig. 5G**, PentaV-FcRα-null; **Movie S3**; **Fig. S5F**), indicating that ECs utilize FcγRIIB to capture pneumococci opsonized by protein antibodies, consistent with our previous findings for pneumococci opsonized with capsule-targeting antibodies (*33*). In contrast, the FcRα-null mice showed a comparable level of KC-captured bacteria as WT mice. We therefore tested the importance of the complement system for CPP-induced KC bacterial capture using mice lacking C3 (the central component of the complement cascade) or the major C3 receptors on mouse KCs CRIg and CR3 (*37*). In C3-deficient mice, KCs showed marked reductions in their ability to capture PentaV immune serum-opsonized pneumococci (**Fig. 5G**, PentaV-C3 KO; **Movie S4**; **Fig. S5F**). Similarly, mice lacking CRIg and CR3 also showed a significantly impaired ability to capture opsonized bacteria (**Fig. 5G**, PentaV-CR KO; **Movie S4**; **Fig. S5F**). Consistent with the deficiency in pathogen capture, C3-and CR-deficient mice were more susceptible to pneumococcal septic infection (**Fig. 5D-F**; **Fig. S5D-E**).

These results indicate that antibodies targeting the CPPs of encapsulated bacteria promote hepatic immunity by activating C3-CR interactions on KCs and, to a lesser extent, IgG-FcγR interactions on ECs. To understand the similarity and differences in the mechanisms of immune clearance of *S. pneumoniae* from the blood after vaccination with protein-or capsule-based vaccines, we characterized the features of PCV13-induced hepatic pathogen capture by IVM. Whilst opsonization with PCV13 immune serum enabled both KCs and ECs to capture blood-borne *S. pneumoniae* in mouse liver sinusoids, the vast majority of immobilized bacteria were associated with ECs and this was dependent on expression of FcRα (**Fig. 5H**, PCV13-FcRα-null). In contrast, KCs were only responsible for a small fraction of the liver-immobilized bacteria, and mice lacking C3 or C3 receptors did not show any obvious defect in PCV13-induced pathogen capture. This result agrees with our previous observation that capsular polysaccharide conjugate vaccines elicit EC-dominant pathogen capture (*33*). In short, although CPP- and capsule-based vaccines induce similar antibody subclass profiles **(Fig. S5G)**, they support markedly different immune clearance mechanisms in liver sinusoids.

### The CPPs are partially localized at or near the external surface of the capsular mesh

To determine how anti-CPP antibodies orchestrate the immune cells and molecules to trap circulating encapsulated bacteria in the liver, we quantified IgG binding and C3 deposition on pneumococci by flow cytometry. PentaV- and PCV13-induced antibodies bound to a comparable proportion of pneumococci, although PCV13 serum produced higher IgG intensity on individual bacteria (**Fig. 6A**). Given the strong dependence of PentaV-mediated bacterial clearance on the complement system, we then assessed C3 deposition triggered by protein-targeting antibodies. Similar to the IgG-binding pattern, PentaV and PCV13 sera generated comparable proportions of C3-positive bacteria (**Fig. 6B**). Consistent with the surface location of the capsule, PCV13 serum induced higher C3 deposition intensity on individual pneumococci than PentaV serum. Similar results were obtained with serotype-6A and -15A pneumococci (**Fig. S6**). These observations indicate that CPPs are accessible to antibodies and that protein-bound antibodies can trigger C3 deposition on intact encapsulated pneumococci.

**Fig. 6.**
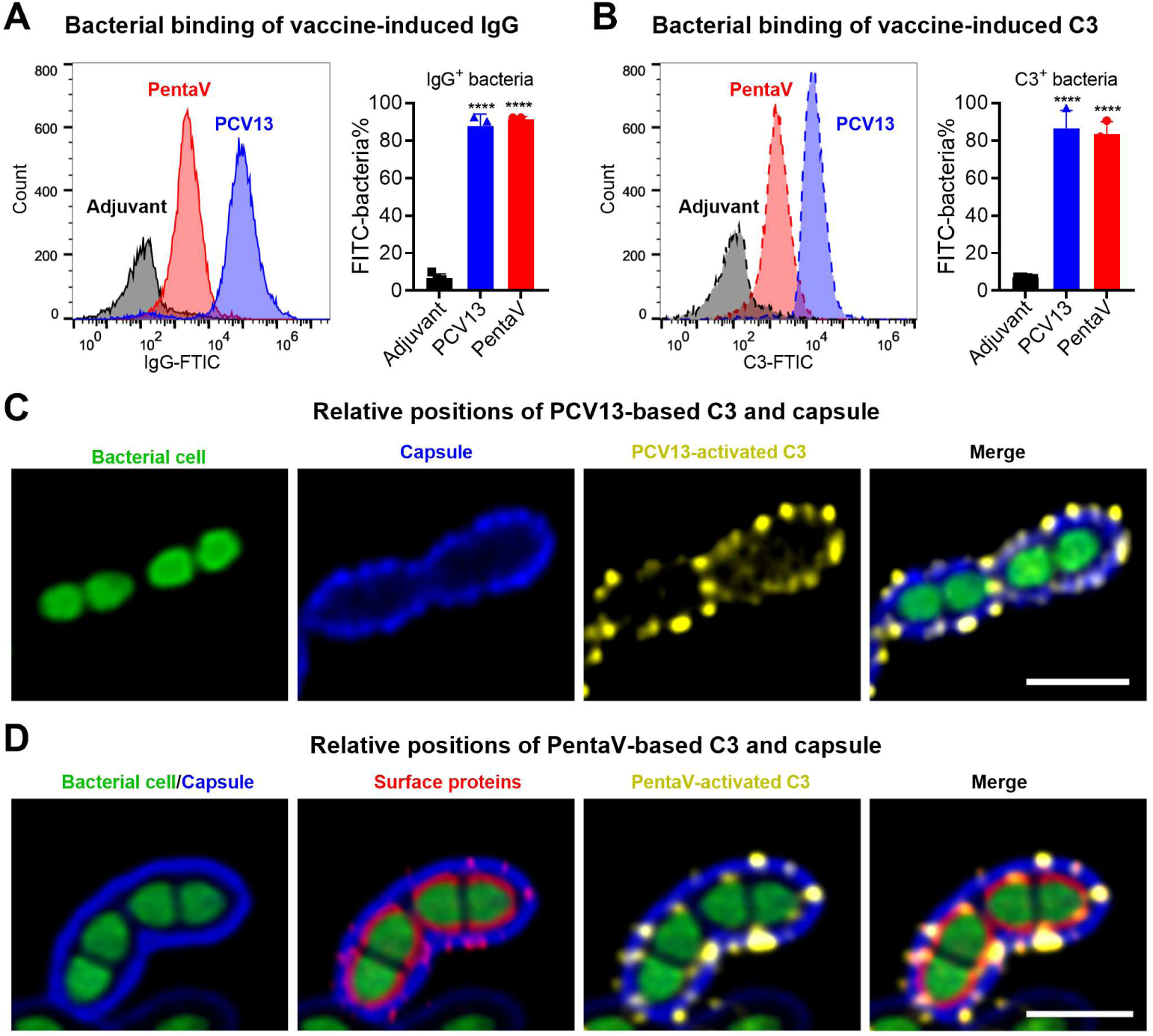
Distinct spatial localizations of protein-attracted antibodies and C3 in the capsular context. **(A)** IgG binding to TIGR4 pneumococci after incubation with mouse anti-protein serum (PentaV) or anti-capsule serum (PCV13) was analyzed by flow cytometry using FITC-conjugated anti-mouse IgG antibody. IgG-bound bacteria are presented as the percentage of total bacteria. n = 3. **(B)** C3 deposition on encapsulated TIGR4 pneumococci was tested and analyzed as in (A), except that bacteria were stained with goat anti-mouse C3 antibody followed by FITC-conjugated anti-goat IgG secondary antibody. n = 3. **(C)** Spatial localization of capsule-associated IgG and C3 deposition on PCV13 serum-opsonized pneumococci. GFP-expressing TIGR4 pneumococci were sequentially incubated with mouse PCV13 antiserum, AF647-conjugated donkey anti-mouse IgG, goat anti-mouse C3 and AF594-conjugated donkey anti-goat IgG antibody. Pneumococci are shown in green, capsule-associated IgG in blue, and C3 in yellow. Signals are pseudocolored as indicated. Scale bar, 2 μm. **(D)** Spatial localization of protein-associated IgG (red), C3 (yellow) and capsule (blue) on PentaV serum-opsonized pneumococci (green). GFP-expressing TIGR4 pneumococci were sequentially incubated with PentaV antiserum, DyLight 405-conjugated donkey anti-mouse IgG antibody, goat anti-mouse C3 IgG, AF594-conjugated donkey anti-goat IgG antibody, heat-inactivated rabbit PCV13 antiserum and AF647-conjugated donkey anti-rabbit IgG antibody. Scale bar, 2 μm. The representative data of at least two independent experiments are presented as mean ± SD, and the statistical differences were determined by multiple *t* tests (A and B). \*\*\*\**P* < 0.0001.

It is generally believed that the capsule reduces antibody recognition of underlying protein antigens (*60–62*). We thus determined the spatial localization of CPP-targeting antibodies on the *S. pneumoniae* surface by confocal microscopy. Staining with capsule antibodies resulted in a uniform fluorescence layer surrounding the bacterium (**Fig. 6C**). By comparison, incubation with PentaV antibodies resulted in focal clusters of immunofluorescence staining within or at the surface of the capsule (**Fig. 6D**), suggesting that at least some of the proteins included in PentaV were able to penetrate the thick capsular layer. However, the PentaV antibody-treated bacteria also showed a nearly continuous immunofluorescence layer beneath the capsule, indicating that significant amounts of the CPPs are localized in the subcapsular space. The presence of abundant antibodies underneath the capsule also revealed that the capsule permits antibodies to access subcapsular proteins, and thus challenges the prevailing view on antibody impermeability of the capsules (*60–62*).

C3 staining revealed distinct spatial patterns of complement activation between pneumococci pre-treated with the protein and capsule antibodies. The PCV13 serum-treated bacteria showed a similar pattern of antibody binding and C3 deposition at the bacterial surfaces and septal regions (**Fig. 6C**). By contrast, the C3 clusters were almost exclusively co-stained with protein antibodies at or near the capsular surface in PentaV serum-treated pneumococci; the abundant antibodies beneath the capsular layer rarely overlapped with C3 (**Fig. 6D**). The strikingly different spatial patterns of protein-targeting antibodies and C3 deposition indicate that the capsule permits antibody access to underlying proteins but restricts the C3 activation/deposition within the deeper subcapsular space. This suggests that IgG bound to protein molecules localized at or near the capsular surface is particularly effective in promoting pathogen capture in the liver sinusoids via complement receptors or FcγRs. Furthermore, these data identify promotion of C3 deposition as a marker for identifying surface proteins that are accessible to IgG binding within or beyond the capsule layer.

### Capsule permeability of proteins shapes their targetability for vaccine immunity

To ascertain the relationship between the capsule permeability of CPPs and their ability to induce pathogen clearance, we selectively characterized the capsule permeability of the CPPs using C3 deposition as a marker. Five CPPs spanning a range of pathogen-clearance activities were selected, including the three most potent antigens (PavA, RrgA and PnrA) and two with relatively weaker activity (PspA and CbpA). Flow cytometry revealed decreasing proportions of C3-positive pneumococci in the order of PavA, RrgA, PnrA, PspA and CbpA after bacteria were pre-treated with the immune sera raised in mice against the corresponding recombinant proteins (**Fig. 7A**). PavA (71.4%), RrgA (59.6%) and PnrA (55.4%) sera yielded the highest proportions of C3-deposited bacteria. This ranking closely mirrored the pattern of pathogen clearance from the bloodstream induced by these proteins (**Fig. 2D**). As the only three sterilizing antigens, PavA, RrgA and PnrA elicited the fastest pathogen clearance with the CT_50_ values of 0.8 min, 1.5 min and 0.9 min (**Fig. 2C; Fig. S2B**). In line with the vaccine-induced bacterial capture and killing, these three proteins also induced the most potent bacterial killing in the liver, with the vast majority of inoculum pneumococci being undetectable at 30 min post i.v. infection (**Fig. 2D**). In agreement with the relatively weaker activities in inducing pathogen clearance, pretreatment with CbpA and PspA sera yielded lower levels of C3-positive pneumococci.

**Fig. 7.**
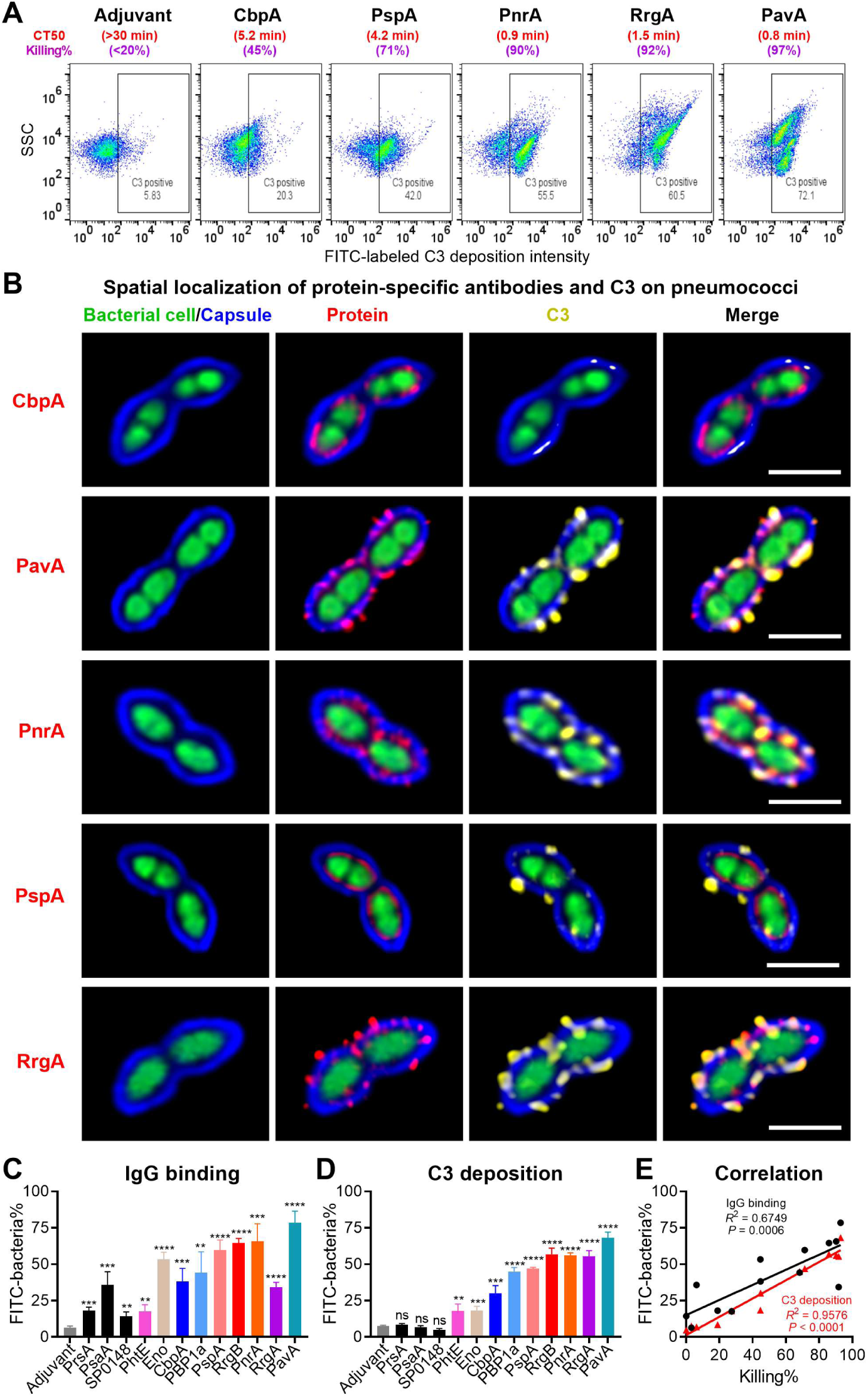
The importance of protein’s capsule permeability in the vaccine targetability. **(A)** C3 deposition on TIGR4 pneumococci was assessed and presented as scatter plot by flow cytometry. The percent C3-labeled bacteria in each panel are indicated with numbers. The relevant CT_50_ (red) value and percent undetectable bacteria (purple) at 30 min post infection are provided from the data in Fig. 2C and Fig. S2C. **(B)** Spatial localization of protein-specific IgG and C3 in the context of the capsule was visualized and presented as in Fig. 6D. Scale bar, 2 μm. **(C** to **E)** The percentage of positive IgG-bound (C) and C3-positive (D) encapsulated TIGR4 pneumococci after incubation with mouse protein immune sera were detected by flow cytometry. IgG- or C3-positive bacteria are presented as the percentages of total bacteria. The correlation between IgG binding or C3 deposition and immune serum-mediated bacterial killing was assessed by linear-regression analysis (E). n = 4-5. The representative data of at least two independent experiments are presented as mean ± SD, and the statistical differences were determined by multiple *t* test (C and D) and *F* test (E). ns, no significant difference; \*\**P* < 0.01; \*\*\**P* < 0.001; \*\*\*\**P* < 0.0001.

We further characterized the surface exposure of these CPPs by confocal immunofluorescence microscopy. Consistent with the flow cytometry data, PavA, RrgA and PnrA were abundantly localized at or near the capsular surface by antibody binding and C3 deposition (**Fig. 7B**). By comparison, CbpA and PspA were predominantly found underneath the capsule, with sporadic distribution across the barrier structure. Additional analyses revealed that RrgB was abundantly distributed at or near the capsular surface, but enolase, PBP1a and PhtE displayed sporadic penetration across the capsular mesh (**Fig. S7**). These spatial patterns were generally in line with their clearance activities against TIGR4 (**Fig. 2D**). This result demonstrated intrinsic variability of CPPs in their spatial localization in the context of the capsular polysaccharide mesh. In addition, C3 deposition, rather than antibody binding intensity, correlated strongly with passive antibody-mediated bacterial clearance (**Fig. 7C-E**), consistent with the importance of antigen proximity to the capsular surface. Collectively, these experiments show that the capsule permeability of extracellular proteins in encapsulated bacteria defines their targetability by vaccine-induced immunity.

## DISCUSSION

The limitations of capsule-based vaccines and the escalating crisis of multidrug resistance underscore the urgent need to develop capsule-independent vaccines against encapsulated bacteria (*1, 63*). Although in pre-clinical studies a large number of extracellular proteins have been studied as vaccine candidates for multiple bacterial pathogens, the only vaccine in clinical use for an encapsulated bacterial pathogen that targets protein antigens is the *N. meningitidis* serogroup B vaccine (*7, 8*). An outstanding challenge in this area is the lack of *in vivo* correlates reflecting how protein antigens enable hosts to clear pathogens in the process of infection. Previous studies have linked the surface accessibility of pneumococcal proteins to antibody-mediated protection (*64*). However, systematic approaches integrating antigen accessibility with functional evaluation of bacterial clearance remain lacking. Based on our recent discovery of the liver-based vaccine mechanisms against encapsulated bacteria (*33, 34*), this study has developed a functional screening approach (L-VAC) to identify capsule-permeable proteins (CPPs) that induce the hepatic clearance of encapsulated bacteria. This approach has enabled us to identify and stratify multiple CPPs in *S. pneumoniae* on the basis of their capability in eliciting hepatic capture of blood-borne bacteria in mice.

The framework of L-VAC is illustrated in **Fig. 8**. It can be summarized in two major steps: 1) enrichment and identification of CPP candidates through antibody binding to intact encapsulated bacteria, with the capsule serving as a physical barrier that restricts antibody access to underlying protein antigens; 2) functional evaluation of antibodies to these proteins for inducing pathogen clearance from the blood circulation (bacterial half-life or CT_50_), including quantification and real-time visualization of protein-elicited pathogen capture in the liver sinusoids of mice (bacterial load in the liver and relative portions of KC/EC-captured bacteria). Thus far, this combination identified IgG targeting seven *S. pneumoniae* proteins that induce significant pathogen clearance, showing the promise of the approach in identifying effective protein antigen vaccine candidates for encapsulated bacteria. The majority of the proteins have been previously characterized, but the L-VAC approach has now allowed the quantitative comparison of pathogen clearance by antibodies to each antigen which is an essential step for the rational selection of antigen candidates for inclusion in future universal pneumococcal vaccines. The L-VAC approach is readily applicable to other bacterial pathogens, and would be suitable for identifying CPPs for other encapsulated bacteria for which there are no existing vaccines in clinical use, including highly antimicrobial-resistant ‘superbugs’ such as *K. pneumoniae* and *Acinetobacter baumannii*.

**Fig. 8.**
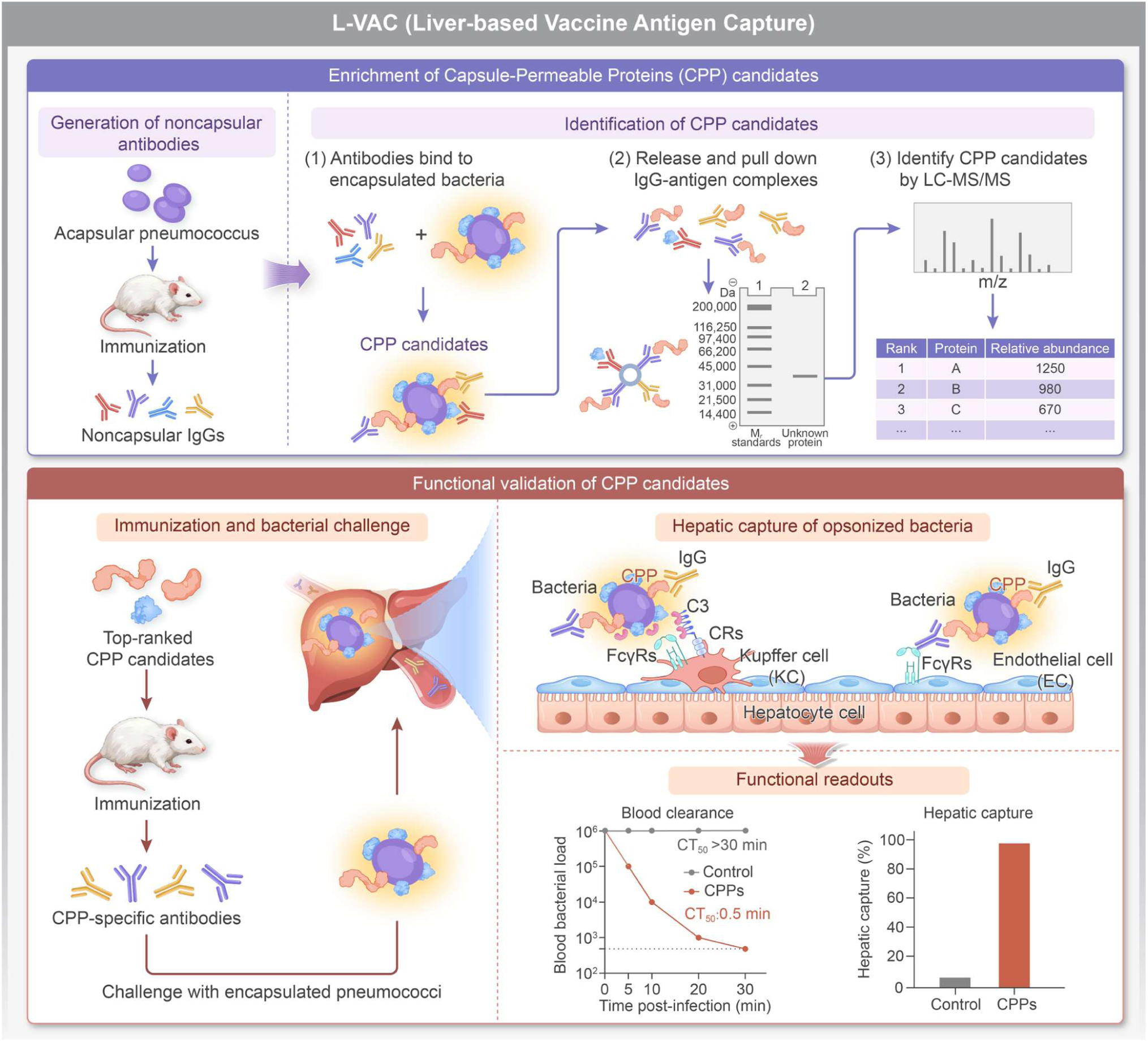
Schematic illustration of the L-VAC platform. The L-VAC strategy consists of two major steps: 1) Enrichment and identification of capsule-permeable proteins (CPPs) candidates through antibody binding to intact encapsulated pneumococci with immune serum raised against acapsular bacteria, followed by isolation of IgG-antigen complexes and LC-MS/MS analysis; 2) Functional validation of CPP candidates by immunization and bacterial challenge, assessing antibody-mediated bloodstream clearance and hepatic bacterial capture by Kupffer cells (KCs) and endothelial cells (ECs). Candidates exhibiting significant pathogen clearance activity are identified as functional CPPs.

Our data have several practical implications for the pre-clinical phase of protein-based vaccine development for encapsulated bacteria. First, antibody binding to noncapsular proteins of encapsulated bacteria cannot be translated into pathogen clearance and host immunity unless it occurs at or near the capsular surface. We found that proteins underlying the capsules are accessible to antibodies, but are unavailable for engaging antibody and complement receptors. Second, while many extracellular bacterial proteins are generally referred to as surface proteins, our microscopy work has revealed that individual proteins vary in their abundance and exposure at the capsular surface, and this is difficult to predict computationally. Third, although individual proteins activate relatively lower levels of pathogen clearance compared with the capsule-targeting vaccines, our data for immunization with multiple CPPs support previous studies showing that vaccination with multiple protein antigens mitigates this functional limitation (*32, 65, 66*). Fourth, vaccination with the same proteins may yield different levels of immunity against different *S. pneumoniae* capsular serotypes even if the proteins are conserved between strains. Similar data have been shown for *A. baumannii* (*67*), and may be caused by capsule type-specific shielding effects on surface accessibility of CPPs and/or variation in surface expression levels of individual antigens. Lastly, this work has further demonstrated that the liver is the major immune organ that executes the protein-induced capture and killing of invasive bacteria, extending our previous data showing the dominance of the liver for executing the capsular polysaccharide vaccine-elicited clearance of pneumococci (*33, 34*). These findings may be used for improving the efficiency in developing broadly protective vaccines against encapsulated bacterial pathogens, particularly the “superbugs” with extensive antimicrobial resistance (*1, 63*).

A striking and important observation is the demonstration of major differences in the mechanisms involved in hepatic clearance of pneumococci between IgG to capsule versus protein antigens. At the cellular level, the protein antigens tested in this work predominantly engage the macrophages and, to a lesser extent, the ECs in the liver sinusoids, whereas the capsular polysaccharide conjugate vaccine (PCV13) predominantly enables the hepatic ECs rather than macrophages to immobilize IgG-opsonized pneumococci (*33*). At the molecular level, the protein antigen-induced bacterial capture predominantly relies on the C3-complement receptor interactions, which is in sharp contrast with the dominant role of the IgG-FcγRIIB recognition in the PCV13-activated pathogen capture (*33*).

The inferiority of the hepatic ECs to macrophages in capturing anti-protein IgG-opsonized bacteria is probably related to the limited exposure of protein antigens at the bacterial surface compared to the abundant surface display of capsular polysaccharides. IgG bound to protein antigens in the subcapsular space seems to remain inaccessible to FcγRIIB, the only Fcγ receptor on the ECs (*33*). Moreover, the relatively sparse protein antigens at bacterial surface may also dampen the “zippering effect” or avidity of the IgG-Fcγ receptor interactions (*33*), and antibody-mediated bacterial agglutination that enhances phagocytic clearance of *S. pneumoniae* (*68*). In contrast, Kupffer cells express multiple Fcγ and complement receptors (*69*), which make the liver macrophages more suitable for recognizing IgG bound to protein antigens. We postulate that the activation of complement C3 by antibodies bound to the capsule-permeable proteins partially overcomes the spatial and abundance “deficits” of protein-bound antibodies by creating a focal mass of potential ligands (C3/iC3b) to allow engagement of multiple complement receptors on the macrophages rather than single IgG-FcγIIB interaction on the ECs. Weaker EC engagement by protein antigens provides a mechanistic explanation for the lower levels of the vaccine-induced immunity for protein antigens compared to capsular polysaccharide (*5*). These mechanistic insights have strongly suggested that the intrinsic “weakness” of the CPPs may be overcome by: 1) immunizing hosts with multiple proteins to maximize target proteins at the capsular surface, and 2) optimizing vaccine formulation to saturate FcγR-reachable target proteins with antibodies.

We have shown that the capsule permeability of protein antigens mainly defines their capability of inducing the hepatic capture of encapsulated bacteria and vaccine potency. The confocal microscopy results revealed that the proteins that elicited sterilizing clearance of TIGR4 pneumococci (e.g., PavA and PnrA) are displayed closer to the bacterial surface than those inducing relatively weaker immunity (e.g., CbpA, PBP1a and PspA). Flow cytometry experiments revealed a different pattern in C3 deposition between the two groups of proteins. This finding should be valuable for selecting which antigens from the large number of protein antigens reported as potential vaccine candidates for *S. pneumoniae* should be taken forward into further preclinical development (*5, 6, 70*). Although the properties defining the capsule permeability of proteins remain to be defined, the protein length does not seem to be a contributing factor since the proteins with superior capsule permeability (e.g., PavA with 551 AAs and PnrA with 350 AAs) are not longer than those with inferior capsule permeability (e.g., CbpA with 693 AAs and PspA with 744 AAs). Moreover, surface attachment mechanisms don’t determine the capsule permeability as the CPPs identified in this work are attached to the phosphorylcholine moiety of teichoic acids (CbpA, PspA), cell wall peptidoglycan (RrgA) or cell membrane (PBP1a, PnrA). However, the significant impact of pneumococcal serotypes on the protein-induced hepatic pathogen clearance does suggest that the chemical and biological properties of different capsular serotypes significantly affect antibody accessibility to underlying proteins.

Taken together, we have developed an *in vivo* platform to functionally identify capsule-permeable protein antigens that induce the hepatic clearance of blood-borne bacteria, the causes of bacterial sepsis. Due to the dominance of drug-resistant encapsulated bacteria in septic infections (*71*), this approach offers a promising strategy to develop broadly protective vaccines against bacterial sepsis.

## MATERIALS AND METHODS

### Bacterial strains and cultivation

All bacterial strains used in this study are listed in **Table S4**. *S. pneumoniae* strains were cultured in Todd-Hewitt broth (Oxoid) supplemented with 0.5% yeast extract (THY) or on tryptic soy agar (TSA, BD Difco) plates supplemented with 5% defibrinated sheep blood at 37°C with 5% CO_2_ as described (*72*). *E*. *coli* and *K. pneumoniae* strains were cultured in Luria-Bertani (LB) broth or on LB agar plates at 37°C under aerobic conditions (*38*).

### Reagents

All chemical reagents were purchased from Sigma-Aldrich (Shanghai, China) unless otherwise noted. DNA processing enzymes were purchased from New England Biolabs (Beijing, China). Primers were synthesized by Xianghong Biotech (Beijing, China) and described in **Table S5.**

### Mice

All mouse experiments were conducted in accordance with protocols approved by the Institutional Animal Care and Use Committee of Tsinghua University. All wild-type mice were purchased from Vital River (Beijing, China). All genetic knockout (KO) mice were generated in the C57BL/6J background. C3 KO mice were acquired from Jackson Laboratory. CRIg/CR3-deficient mice were generated in our previous study (*33*). FcRα-null mice lacking all FcγR α-chain genes and FcγRs-humanized mice were generated as described (*73*) and provided by Jeffrey V. Ravetch and Fubin Li. *Clec4f*-DTR mice were provided by Martin Guilliams (*74*). KCs were depleted in *Clec4f*-DTR mice by i.v. injection of diphtheria toxin (25 ng/g body weight) 12 h before infection. All mice were kept under specific pathogen-free (SPF) conditions with free access to food and water.

### Bacterial mutagenesis

The acapsular pneumococcal strain TH7660 was generated from TH1759, a streptomycin-resistant derivative of the serotype-4 strain TIGR4, by replacing the capsule biosynthesis locus using Janus Cassette-mediated counterselection as described (*75*). GFP-expressing pneumococci were generated by inserting a GFP-kan cassette downstream of the *aliA* gene as described (*39*). The capsule- and O-antigen-deficient *K. pneumoniae* strain TH16938 (ATCC43816 background, K2) was generated from the acapsular strain TH13811 by deleting O-antigen biosynthesis genes using a CRISPR-based mutagenesis approach as previously described (*38*). Briefly, the *rfb* gene cluster was deleted using flanking regions amplified with primer pairs Pr19323-Pr19326.

### Immunogen preparation

Inactivated acapsular bacteria used as whole-cell vaccines (WCVs) were prepared by treating freshly cultured acapsular *S. pneumoniae* TH7660 and *K. pneumoniae* TH16938 with 0.5% formaldehyde overnight at 4°C as described (*33*). Protein antigens were prepared by amplifying the coding sequences of target proteins from the TIGR4 genome, and cloning them into the NcoI/BamHI sites of pET28a with an N-terminal Strep II tag (*76*), or downstream of the glutathione S-transferase (GST) gene in the NotI/KpnI sites of pGEX-2T (*31*). An HRV 3C protease recognition site (LEVLFQGP) was introduced between the GST and the target protein in pGEX-2T to enable tag removal during purification. The inserts of all recombinant plasmids were confirmed by DNA sequencing. The GST tag was cleaved with PreScission Protease (Beyotime, P2303) according to the manufacturer’s instructions. Purified proteins were analyzed by SDS-PAGE, concentrated using centrifugal filters (Millipore, UFC903008) in phosphate-buffered saline (PBS) and quantified by the bicinchoninic acid (BCA) assay kit (Beyotime, P0012S). All relevant primers and plasmids are shown in **Tables S5** and **S6**.

### Immunoprotection in mice

Immunization was carried out in 6-8-week-old CD1 mice as described previously (*31, 33*). Briefly, mice were subcutaneously immunized with inactivated acapsular pneumococci (5 × 10^8^ CFU in 50 μl PBS), individual recombinant proteins (5 µg in 50 μl PBS), or the pentavalent protein vaccine (50 µg in 50 μl PBS), each emulsified with 50 μl Freund’s complete adjuvant (Sigma, F5881). Mice were boosted twice at a 14-day interval with the same dose without adjuvant. PCV13 immunization was carried out as described (*33*).

Vaccine-induced protection was assessed by measuring hepatic clearance of blood-borne bacteria during the first 30 min after i.v. infection, and by monitoring survival for 7 days or until mice reached a humane endpoint defined as > 20% body weight loss. Bacteremia kinetics were assessed by CFU plating as described (*37*). The time for 50% bacterial clearance from the blood circulation (CT_50_) was calculated from bacteremia kinetics using one-phase association nonlinear regression as described (*37*). Total bacterial burden in each animal was calculated as the sum of CFU recovered from blood and homogenates of the major organs.

Passive protection against bacterial blood infection was evaluated as described (*33*). Immune serum or purified IgG was pre-incubated with bacteria in 100 μl PBS for 10 min at room temperature (RT) before i.v. injection into naïve mice. Unless otherwise indicated, serum samples were heat-inactivated at 56°C for 30 min before being used for bacterial opsonization.

### Flow cytometry

IgG binding and C3 deposition on bacteria were assessed by flow cytometry using immune serum without heat inactivation as described (*77*). Briefly, 10^6^ CFU of pneumococci were incubated in 100 μl PBS containing immune serum raised with protein antigens (20%), acapsular WCV (10%) or PCV13 (10%) at RT for 15 min. After washing with PBS twice, bacteria were incubated with FITC-conjugated goat anti-mouse IgG antibody (1:500; EasyBio, BE0111-100) for 10 min at RT. C3 deposition was detected by further incubating the immune serum-opsonized bacteria with goat anti-mouse C3 antibody (0.5 μg/ml; MP Biomedicals, 55557) for 10 min at RT, and finally with a FITC-conjugated rabbit anti-goat IgG antibody (1:500; EasyBio, BE0120-100) for 10 min at RT. FITC signals were acquired by a flow cytometer (FongCyte^TM^, A24000085) and analyzed using FlowJo (10.10).

### ELISA

Serum antibodies were quantified by enzyme-linked immunosorbent assay (ELISA) as described (*31*). HRP-conjugated mouse anti-human IgG-and IgM-specific antibodies or goat anti-mouse IgM-, IgG-, IgG1-, IgG2a-, IgG2b-, IgG2c-and IgG3-specific antibodies were used to determine the antibody levels and subclasses. Endpoint antibody titers of immune serum were calculated after subtraction of background signals from the adjuvant control group.

### Serum antibody purification and depletion

IgG was purified from murine serum using protein-G resins (GenScript, L00209) according to the manufacturer’s instructions. Purified IgG fractions were assessed by SDS-PAGE, concentrated using centrifugal filters (Millipore, UFC903008), and quantified using a BCA Assay Kit (Beyotime, P0012S). The flow-through fractions were used as IgG-depleted sera. Naturally acquired antibodies against serotype-4 and -6A capsular polysaccharides in normal human sera were depleted using CPS-conjugated beads as described previously (*78*). Briefly, free CPSs were activated with 4-(4,6-dimethoxy-1,3,5-triazin-2-yl)-4-methyl morpholinium chloride (DMTMM; Sigma, 74104-1G-F), ultrafiltered with centrifugal filters (Millipore, UFC801008) to remove the free activator and mixed with carboxyl latex beads (2 μm; Invitrogen, C37278) overnight at RT. The conjugated beads were blocked with 1% bovine serum albumin in PBS. Human sera were incubated with CPS-conjugated beads on a rotor for 2 h at RT. The depletion effect was verified by ELISA.

### Immunoblotting

Antibody binding to encapsulated bacteria was assessed by immunoblotting as described (*76*). Briefly, 1 × 10^8^ CFU of fresh *S. pneumoniae* were washed twice with PBS and incubated with 10 μl immune serum at RT in a total volume of 100 μl for 10 min. After washing with PBS containing 0.05% Tween 20 (PBST) twice, bacterial pellets were resuspended in 100 μl SDS-PAGE reducing buffer and boiled at 96℃ for 15 min for SDS-PAGE and immunoblotting analyses. HRP-conjugated goat anti-mouse IgG antibody (EasyBio, BE0102) was used at a dilution of 1:5000.

### L-VAC screening

Pneumococcal CPPs were identified using the liver-based vaccine antigen capture (L-VAC), adapted from a previously described immunoprecipitation method (*79*). Encapsulated pneumococci of strain TIGR4 were incubated with murine WCV immune serum. Antibody binding to encapsulated bacteria was first assessed by immunoblotting to optimize the experimental conditions, including serum volume and bacterial input. The ratio of bait antibody to total bacterial lysate protein was adjusted to approximately 1:120 -1:150 as described (*80*). Briefly, fresh bacterial culture in THY at OD_620 nm_ = 0.6 was harvested by centrifugation, washed three times with Tris buffered saline (TBS) and resuspended in TBS supplemented with anti-WCV immune serum. After incubation at RT for 30 min, bacteria were pelleted, washed twice in TBS, and resuspended in cell lysis buffer (TBS, pH 7.6, 1 mM PMSF, 10 μg/ml DNase, 20 μg/ml RNase, 200 μg/ml lysozyme, 5% glycerol, 1% Triton X-100), and processed using Pressure Cell Disruptor (ZHICHU, ZQZY-C). The lysate was centrifuged at 10,000 × *g* for 15 min; the supernatants were used to immunoprecipitate the protein-antibody complexes using Pierce^TM^ Protein A/G Magnetic Beads according to the manufacturer’s instructions (Thermo, 88802). Following SDS-PAGE separation and silver staining (Beyotime, P0017S), the resulting proteins were eluted from polyacrylamide gels and analyzed by LC-MS/MS using a Thermo-Dionex Ultimate 3000 HPLC system coupled with a Thermo Orbitrap Fusion mass spectrometer. The acquired MS/MS spectra were queried against the TIGR4 database using Proteome Discoverer software (v1.4). The serum from adjuvant-treated mice was processed in the same manner as a negative control. Antibody-mediated protein enrichment was calculated by comparing the levels of the same proteins between the immune and adjuvant serum groups on the basis of the proteomic data.

### Intravital microscopy (IVM) of the murine liver

IVM imaging of murine liver sinusoids was performed using inverted confocal laser scanning microscopy as described (*37*). Briefly, KCs and ECs were labeled by i.v. injection of AF647 anti-F4/80 and AF594 anti-CD31 antibodies, respectively, before i.v. injection of 5 × 10^7^ CFU FITC-labelled bacteria. Images of the liver sinusoids were acquired under three laser excitation wavelengths (488, 585, and 635 nm) using a white-light laser (1.5 mW, Laser kit WLL2, 470-670 nm) on Leica TCS SP8 confocal microscope (20 × /0.80 NA HC PL APO objective). The LAS X Core module (version 3.7.3) was used to immediately acquire real-time images for at least 3 min after i.v. infection (600 × 600 pixels). At least 5-12 random fields of view (FOV) between 10 and 20 min after infection were chosen to quantify hepatic cell-captured bacteria (1024 × 1024 pixels). For IVM detection of serum-mediated passive clearance, 5 × 10^7^ CFU of FITC-labeled bacteria were pre-incubated with 100 μl heat-inactivated serum for 10 min at RT before i.v. injection to naïve mice.

### Bacterial immunofluorescence microscopy

Pneumococcal capsule, protein antigens, bacterium-bound antibodies and deposited C3 were visualized by immunofluorescence microscopy as described with minor modifications (*81*). GFP-expressing pneumococci (2 × 10^7^ CFU) were incubated with 100% murine protein immune serum or 30% murine PCV13 immune serum, in a total volume of 100 μl for 10 min at RT. After bacteria were washed once with 500 μl PBS by centrifugation and resuspension, the pellets were resuspended in 100 μl PBS containing DyLight 405 donkey anti-mouse IgG antibody (1:100; YEASEN, 34104ES60), and incubated for 10 min at RT. C3 deposition on the bacteria was detected by sequentially treating the abovementioned pneumococci with 20 μl PBS containing goat anti-mouse C3 antibody (1:10; MP Biomedicals, 55557) and 100 μl PBS containing AF594 donkey anti-goat IgG antibody (1:100; YEASEN, 34312ES60) each for 10 min at RT. Bacteria were washed once with 500 μl PBS prior to each treatment. For samples treated with protein immune serum, capsule staining was performed after C3 staining using 30% heat-inactivated rabbit PCV13 immune serum followed by AF647-conjugated donkey anti-rabbit IgG antibody (1:100, PBM, DRB2005).

For microscopic viewing, bacterial pellets were resuspended in 10-15 μl PBS, dropped onto microscope slides (26 × 76 × 1.0 mm) and covered with coverslips (20 mm × 70 mm). Samples were observed with continuous spectrum laser confocal microscopy Leica STELLARIS 8 Falcon (A23000054). Images were acquired using a 405-nm solid-state laser and a white-light laser (440 -790 nm) set to excitation wavelengths of 488, 561 and 638 nm, with a 100 × oil-immersion objective, a 6 × zoom factor, an image size of 1024 × 1024 pixels and a scan speed setting of 200.

### Sequence analysis

DNA and protein sequences were analyzed using SnapGene (4.3.6). Protein sequence identity was determined using BLAST. The CLUSTALW program (https://www.genome.jp/tools-bin/clustalw) was used to compare PspA and CbpA protein sequences. The amino acid sequences of nine proteins from the serotype 4 *S*. *pneumoniae* strain TIGR4 were used as references to search 34,099 pneumococcal genomes in PubMLST (https://pubmlst.org). Query lengths were calculated from the supplied sequences, excluding terminal stop symbols. For each isolate and protein, the reported best-match amino acid identity and alignment length were analyzed. An alignment coverage proxy was calculated by dividing alignment length by the corresponding TIGR4 query length. Matches with a coverage proxy ≥90% were retained without applying a minimum identity threshold. Among these matches, amino acid identity relative to TIGR4 was summarized as the median and range (minimum–maximum). The proportions of retained matches with identity ≥70%, ≥80%, and ≥90% were also calculated. Because alignment length can include gaps, the coverage proxy does not represent exact query coverage, and identity values describe aligned regions rather than independently verified full-length homologs.

### Data analysis

All experiments were carried out with at least two biological replicates. Data are expressed as mean ± standard deviation (SD). Comparisons between two groups were performed using two-sided unpaired *t* tests. Two-way ANOVA with Tukey’s multiple comparisons test and multiple *t* tests were applied to compare more than two groups. Mouse survival data were analyzed by log-rank (Mantel-Cox) test. Flow cytometry data were analyzed using FlowJo (10.10). Linear regression was performed using Pearson’s method and *F* test. These statistical analyses above were performed using GraphPad Prism software (8.0.1). *P* < 0.05 was considered statistically significant (\**P* < 0.05, \*\**P* < 0.01, \*\*\**P* < 0.001, and \*\*\*\**P* < 0.0001).

## List of Supplementary Materials

**Figs. S1 to S7**

**Tables S1.** Mass spectrometry data of bacterial protein antigens by L-VAC

**Tables S2.** Shortlists of bacterial protein antigens for functional evaluation

**Tables S3.** Sequences and identities of capsule permeable proteins among pneumococcal strains

**Tables S4.** Bacterial strains used in this study

**Tables S5.** Primers used in this study

**Tables S6.** Plasmids used in this study

**Movies S1. Noncapsular antigen-activated capture of serotype-4 pneumococci in liver sinusoids.** Liver sinusoid endothelial cells (ECs, cyan) and resident macrophage Kupffer cells (KCs, red) capture of i.v. inoculated serotype-4 pneumococci (5 × 10^7^ CFU, green) from the bloodstream of adjuvant, PentaV- and WCV-immunized mice. Intravital microscopic visualization was initiated immediately before bacterial injection; imaging data at the onset of bacterial appearance in the circulation are presented. The playback rate is eight frames per second. Quantitative data and selected images are shown in Fig. 5A.

**Movies S2. The role of Kupffer cell in PentaV serum-activated capture of serotype-4 pneumococci in liver sinusoids.** Visualization of EC- and KC-associated pneumococci in liver sinusoids of naïve mice with or without DT treatment were imaged as in Movie S1 after i.v. injection of 5 × 10^7^ CFU of PentaV serum-opsonized serotype-4 pneumococci. The bacteria were pre-incubated with 100 µl of adjuvant or PentaV serum for 10 min at RT before injection. Quantitative data and selected images are shown in Fig. 5B.

**Movies S3. The role of Fcγ receptors in PentaV serum-activated capture of serotype-4 pneumococci in liver sinusoids.** Visualization of EC- and KC-associated pneumococci in liver sinusoids of naïve WT and FcRα-null mice were imaged as in Movie S1 after i.v. injection of 5 × 10^7^ CFU of PentaV serum-opsonized serotype-4 pneumococci. Quantitative data and selected images are shown in Fig. 5C.

**Movies S4. The importance of complement C3 / C3 receptor in PentaV serum-activated capture of serotype-4 pneumococci in liver sinusoids.** Visualization of EC- and KC-associated pneumococci in liver sinusoids of naïve WT, C3 KO and CR KO mice were imaged as in Movie S1 after i.v. injection of 5 × 10^7^ CFU of PentaV serum-opsonized serotype-4 pneumococci. Quantitative data and selected images are shown in Fig. 5C.

## ACKNOWLEDGMENTS

We would be grateful to Jeffrey V. Ravetch and Fubin Li for generously providing FcRα-null and FcγRs-humanized mice, and Martin Guilliams for *Clec4f*-DTR mice. This work received technical support from the Tsinghua research platforms in animal experimentation (Laboratory Animal Research Center), flow cytometry (Medical Microbiology Core) and IVM imaging (Center for Cell Biology) and protein mass spectrometry (Center for Proteomics).

## Funding

National Key Research and Development Program of China grants 2023YFC2306300

National Natural Science Foundation of China 82330071

Tsinghua University Initiative Scientific Research Program 20243080033

Conducting Surveys, Research and Risk Assessment on Zoonotic Diseases 0733-24095348

## Author contributions

Conceptualization: QJ, JJW, JRZ

Experimentation: QJ, YRW

Methodology: QJ, YRW, JJW, HTD, JSB

Investigation: QJ, JJW, JRZ

Visualization: QJ, YRW

Funding acquisition: JRZ, WWZ

Project administration: JRZ

Supervision: QJ, JJW, JRZ

Writing – original draft: QJ, JJW, JRZ

Writing – review & editing: QJ, YRW, JJW, JRZ, JSB

## Data and materials availability

All of the data reported in this publication are available upon request.

## SFIGURE AND LEGENDS

**Fig. S1.**
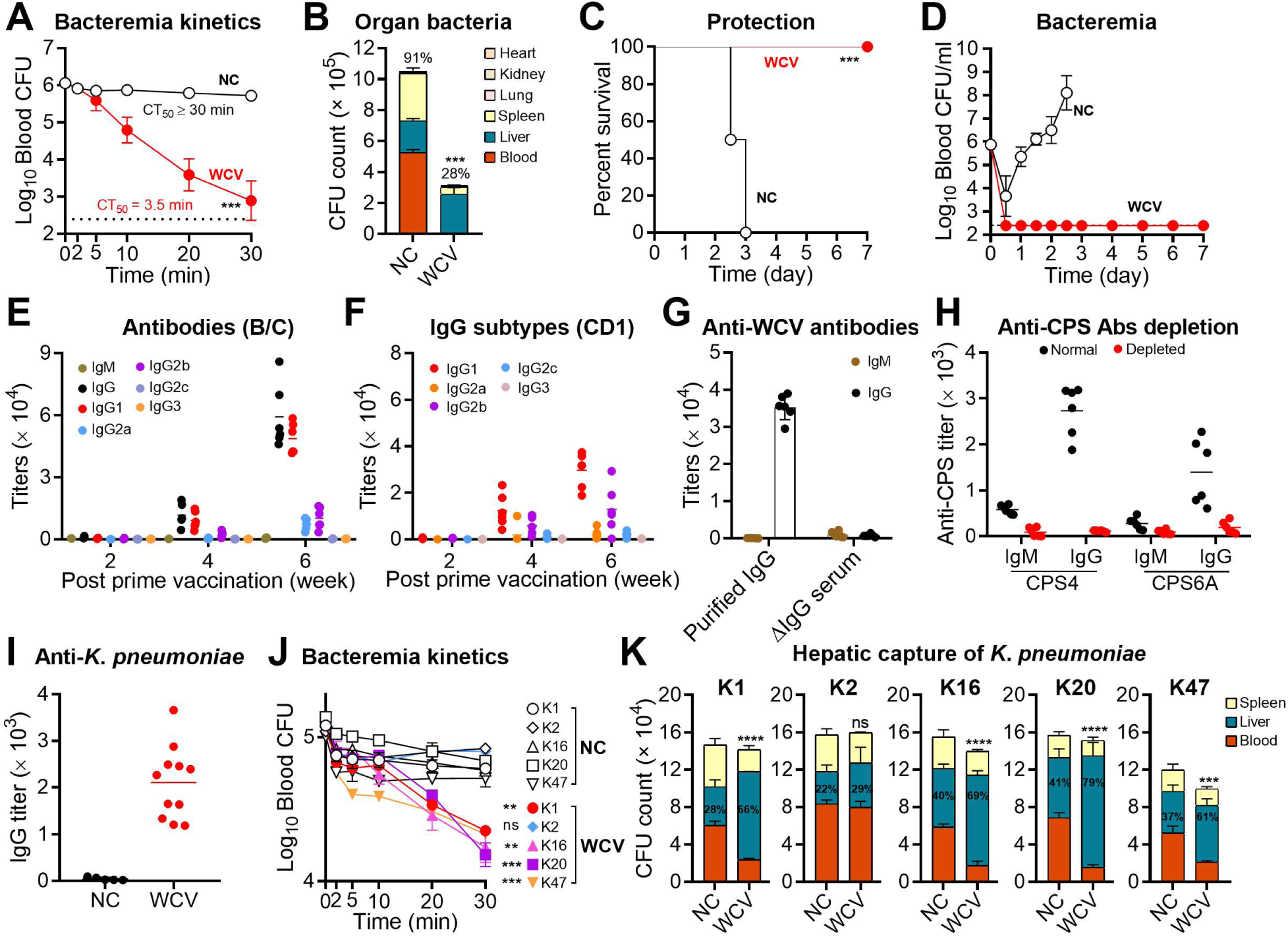
Non-capsular antigens-induced immunity against encapsulated pneumococci in BALB/c mice. **(A** to **D)** Capsule-null WCV immunization promotes protection against encapsulated pneumococci. Bacteremia kinetics (A), bacterial distribution in major organs at 30 min (B), survival (C) and bacteremia levels (D) of WCV- or adjuvant-immunized mice after i.v. infection with 10^6^ CFU of serotype-4 pneumococci. NC, adjuvant control. n =3 (A and B) or 6 (C and D). **(E)** Antibody types against non-capsulated pneumococci in serum of BALB/c mice were measured by ELISA at weeks 2, 4 and 6 after the prime immunization. n = 6. **(F)** Serum IgG subclasses in serum of CD1 mice were measured by ELISA at weeks 2, 4 and 6 after the prime immunization. n = 6. **(G)** IgG and IgM levels in purified IgG fractions from WCV immune serum and in IgG-depleted WCV immune serum (ΔIgG serum) generated using Protein G beads. WCV immune serum was from CD1 mice at week 6 after the prime immunization. n = 6. **(H)** IgG and IgM titers against CPS4 and CPS6A in normal human sera before and after depletion of CPS-specific antibodies using a mixture of CPS4- and CPS6A-conjugated beads. n = 6. **(I)** IgG titers against the capsule/O antigen-deficient *K. pneumoniae* strain in adjuvant control (NC) and WCV immune sera. CD1 mice were subcutaneously immunized twice, two weeks apart, with 5 × 10^8^ CFU of formaldehyde-inactivated CPS/O antigen-deficient *K. pneumoniae* derived from K2 strain ATCC43816 (WCV). Sera were collected from adjuvant control (NC)- or WCV-immunized mice two weeks after the final immunization, and IgG titers were measured by ELISA. n = 5-11. **(J** and **K)** Cross protection against multiple *K. pneumoniae* capsule types mediated by WCV immune serum. Bacteremia kinetics (J) and organ bacteria at 30 min (K) were detected after i.v. infection with 1 × 10^5^ CFU of *Kpn* strains representing capsule types K1, K2, K16, K20, and K47. Bacteria were pre-incubated with 100 μl adjuvant control serum (NC) or WCV immune serum before injection into C57BL/6J mice. Numbers on the column in (K) represent the mean percentage of bacteria trapped in the liver. n = 3. The representative data of at least two independent experiments are presented as mean ± SD, and the statistical differences were determined by two-way ANOVA with Tukey’s multiple comparisons test (A and J), unpaired *t* tests (B and K) and log-rank (Mantel-Cox) test (C). ns, no significant difference; \*\**P* < 0.01; \*\*\**P* < 0.001; \*\*\*\**P* < 0.0001.

**Fig. S2.**
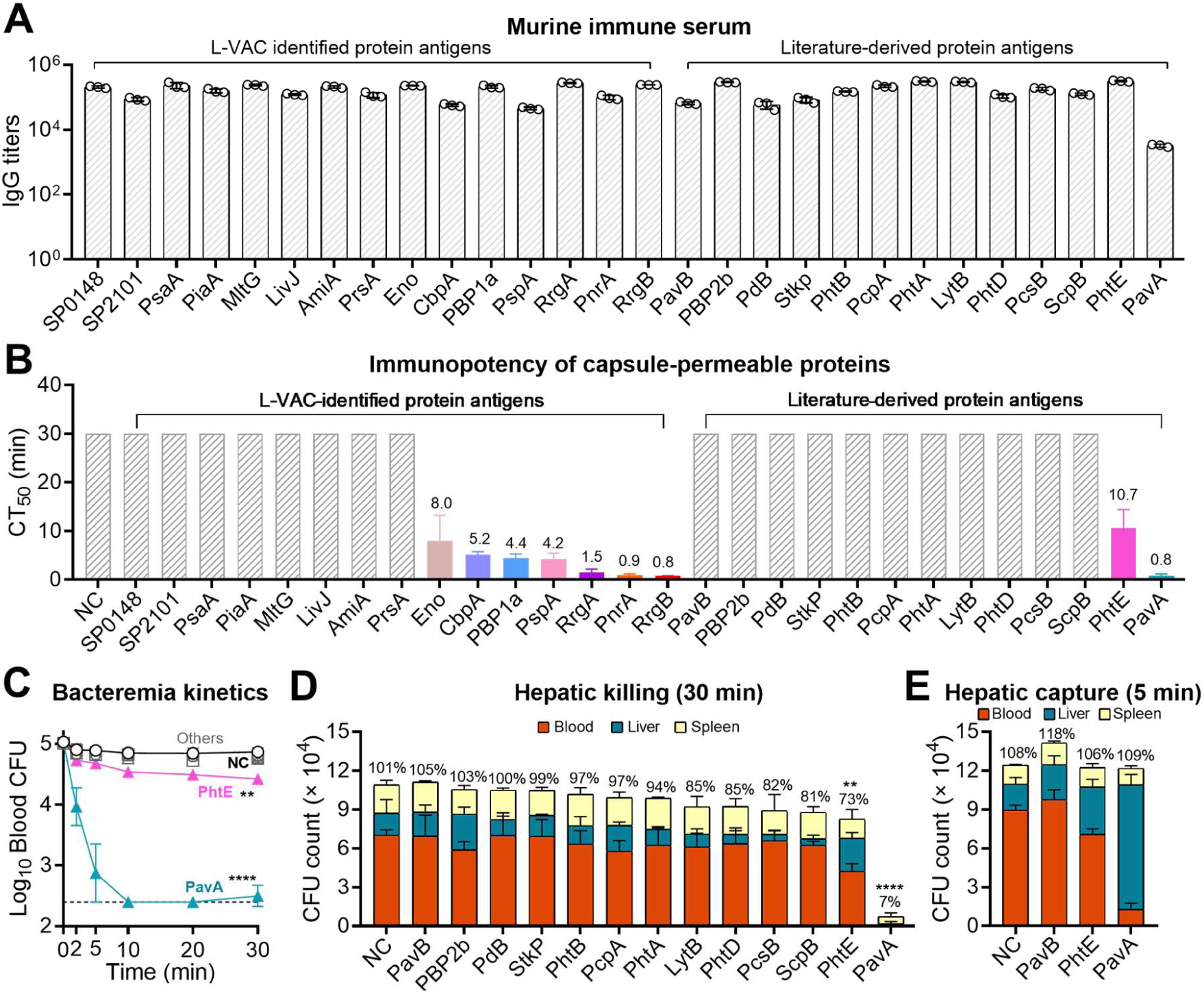
Anti-protein immune serum-triggered hepatic clearance of serotype-4 pneumococci. **(A)** IgG titers against 15 L-VAC candidate proteins and 13 previously reported pneumococcal protein antigens in murine immune sera were detected by ELISA. n = 3. **(B)** Protective activity of candidate proteins in the passive bloodstream clearance model was expressed as the value of CT_50_. NC, adjuvant control. n = 3-5. **(C** to **E)** Protective activity of 13 previously reported pneumococcal protein antigens was evaluated using sera from mice immunized with recombinant proteins. Encapsulated serotype-4 pneumococci TIGR4 were pre-opsonized with 20% immune sera and i.v. inoculated into naïve CD1 mice to assess bacterial clearance from the bloodstream (C), bacterial killing (D) and bacterial capture (E) in the liver. NC, adjuvant control. n = 3-6. The representative data of at least two independent experiments are presented as mean ± SD, with statistical differences determined by two-way ANOVA with Tukey’s multiple comparisons test (C) and multiple *t* tests (D). \*\**P* < 0.01; \*\*\*\**P* < 0.0001.

**Fig. S3.**
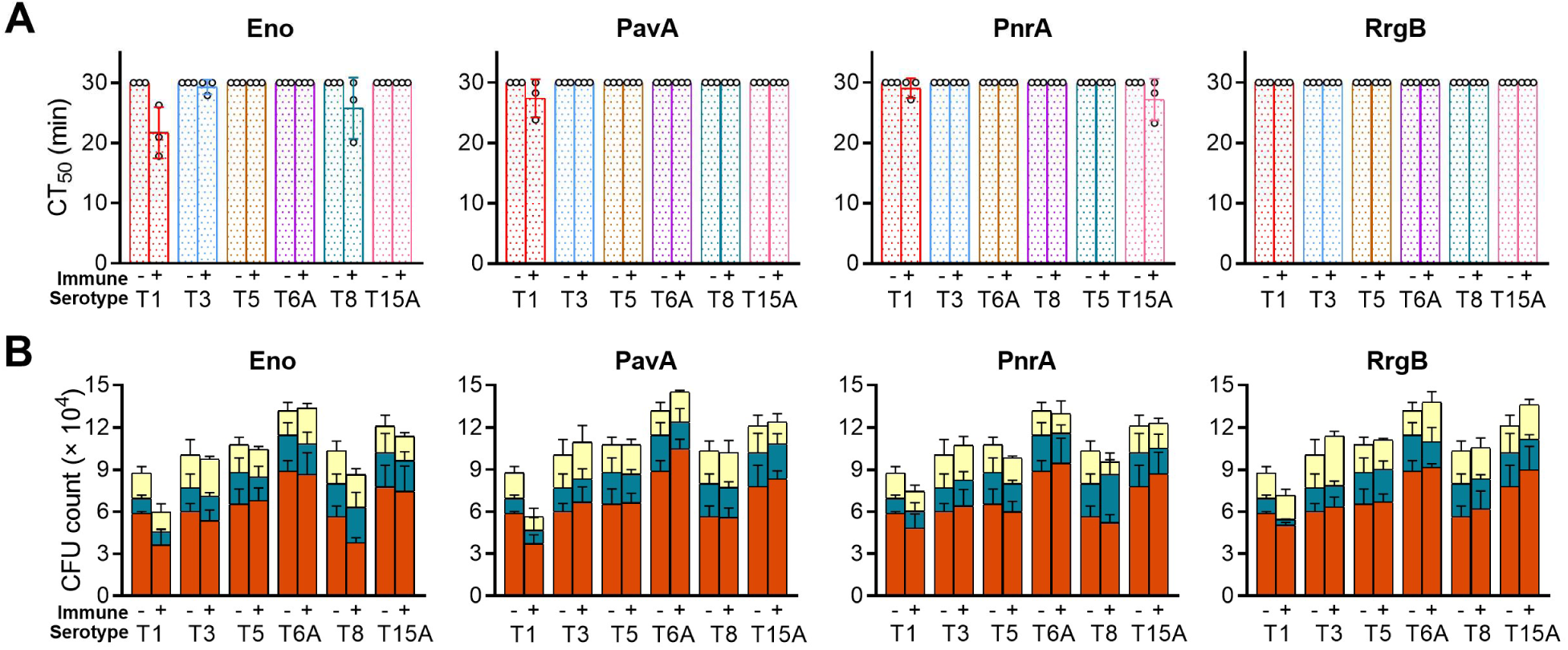
Cross-serotype protective activity of additional pneumococcal proteins. **(A** and **B)** Cross-serotype protective activities of Eno, PnrA, PavA, and RrgB were evaluated using sera from mice immunized with recombinant proteins. Serotypes 1, 3, 5, 6A, 8, and 15A of *S. pneumoniae* were pre-opsonized with 20% immune serum and i.v. inoculated into naïve CD1 mice at 10^5^ CFU per mouse to assess bacterial clearance from the bloodstream (A) and organ bacteria at 30 min (B). -, adjuvant control; +, protein immune serum. n = 3. The representative data of at least two independent experiments are presented as mean ± SD.

**Fig. S4.**
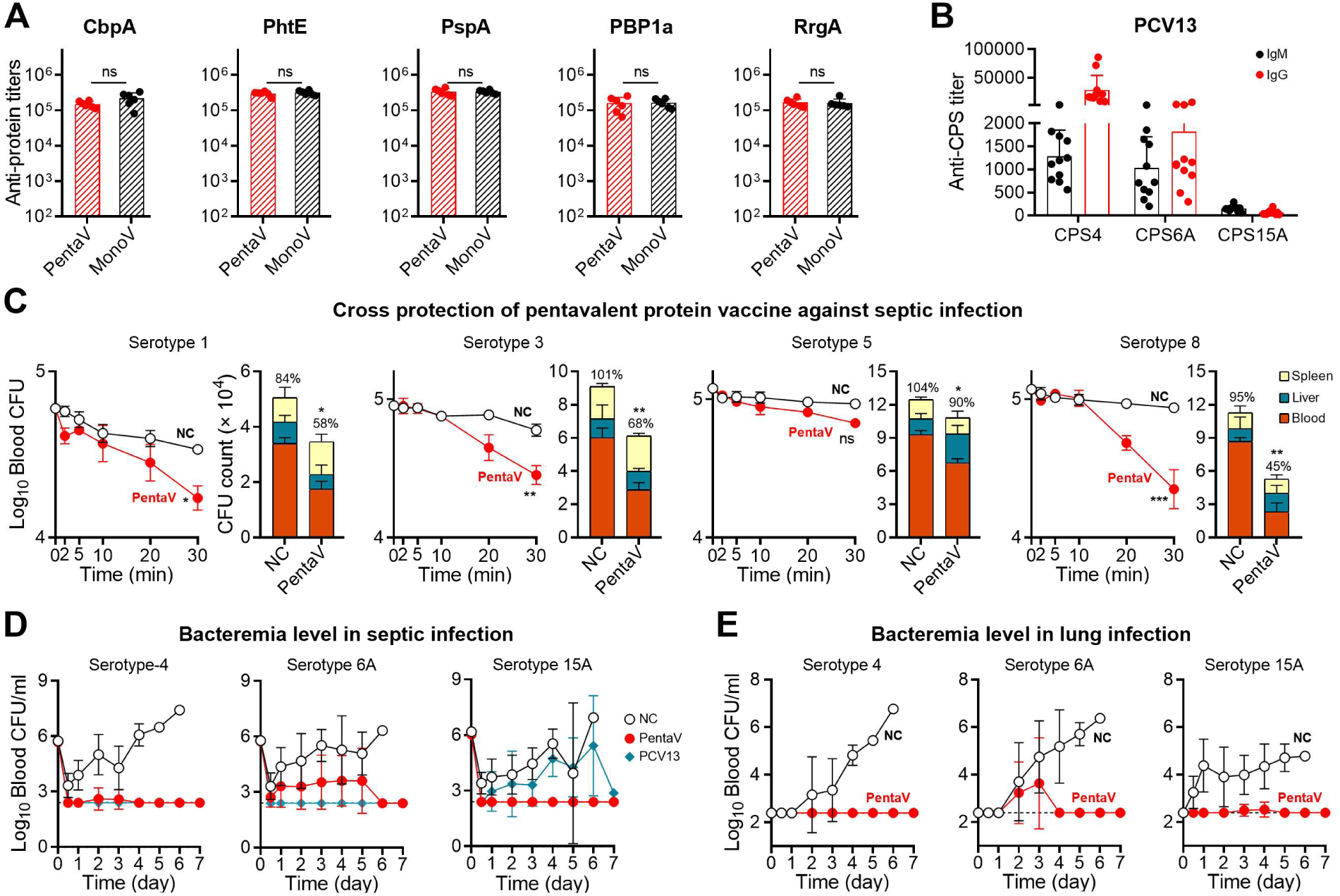
Cross-serotype immunity conferred by the pentavalent protein vaccine. **(A)** IgG titers against PhtE, CbpA, PBP1a, PspA and RrgA in sera from mice immunized with the pentavalent (PentaV) and the corresponding monovalent protein vaccine (MonoV) were detected by ELISA. n = 6. **(B)** IgM and IgG titers against CPS4, CPS6A and CPS15A in PCV13-immunized sera were detected by ELISA. n = 11. **(C)** Cross-serotype protection conferred by PentaV against multiple pneumococcal serotypes. Bacteremia kinetics during the first 30 min (left panel) and organ bacteria at 30 min (right panel) were determined after i.v. infection of adjuvant control (NC)- and PentaV-immunized mice with 1 × 10^5^ CFU of serotype*-*1, -3, -5, and -8 pneumococci. n = 3. **(D)** Bacteremia levels of adjuvant control (NC)-, PentaV- and PCV13-immunized mice over 7 days after intravenous septic infection with 10^6^ CFU of serotype-4 or -6A pneumococci, or 2 × 10^6^ CFU of serotype-15A pneumococci. n = 6-11. **(E)** Bacteremia levels of adjuvant control (NC)- and PentaV-immunized mice over 7 days after intranasal lung infection with 2 × 10^6^ CFU of serotype-4 pneumococci, 10^6^ CFU of serotype-6A pneumococci, or 2 × 10^7^ CFU of serotype-15A pneumococci. n = 6. The representative data of at least two independent experiments are presented as mean ± SD, with statistical differences determined by unpaired *t* test (A), two-way ANOVA with Tukey’s multiple comparisons test (left panel in D) and multiple *t* tests (right panel in D). ns, no significant difference; \**P* < 0.05; \*\**P* < 0.01; \*\*\**P* < 0.001.

**Fig. S5.**
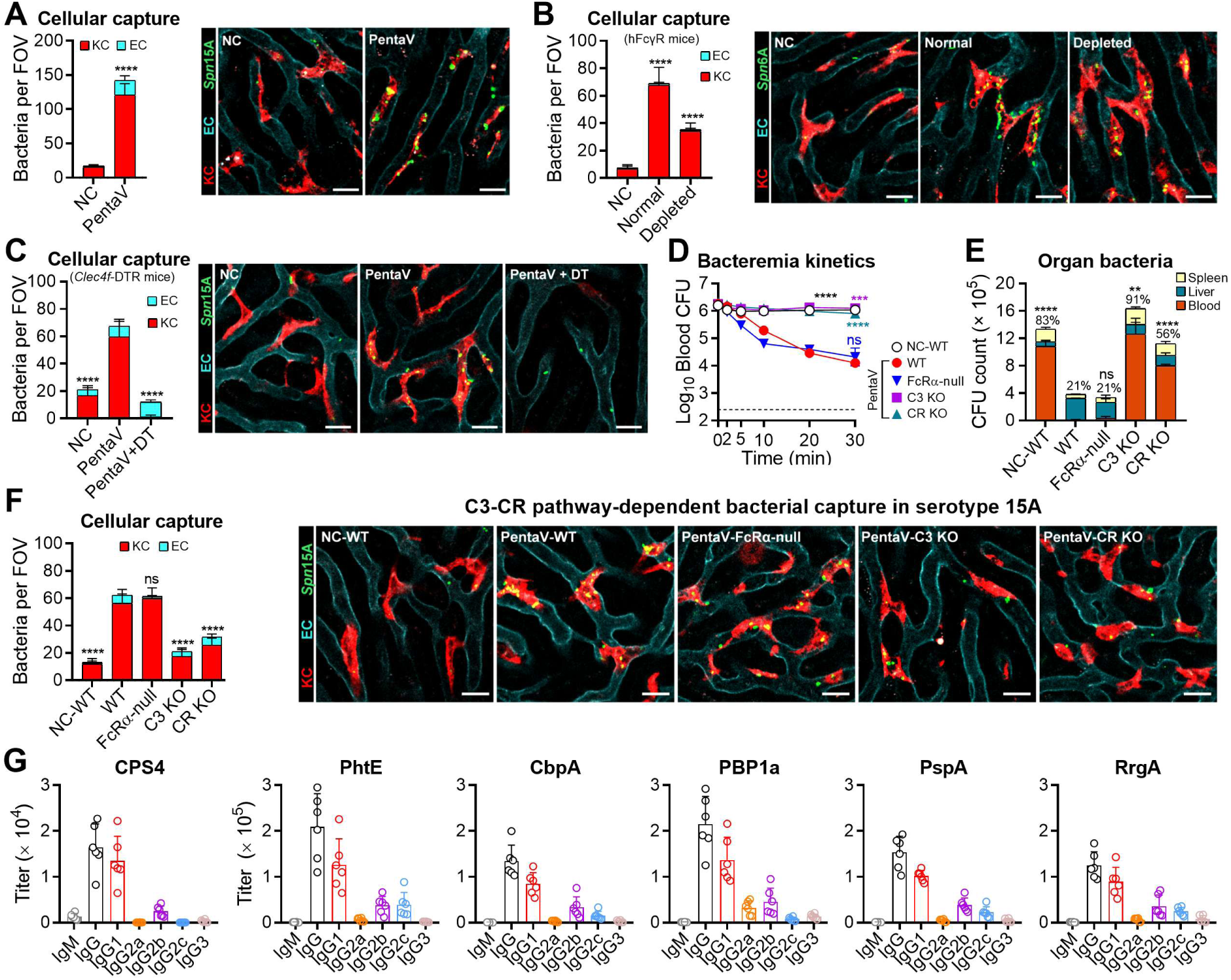
The roles of liver macrophages and the complement system in protein-induced hepatic clearance of blood-borne pneumococci. **(A)** Representative IVM images and quantitation of immobilized serotype-15A pneumococci in PentaV-immunized mice as in Fig. 5. Scale bar, 20 μm. n = 6-12. **(B)** Same as in (A) with the exception of using serotype-6A pneumococci for infection. Scale bar, 20 μm. n = 6-12. **(C)** The impact of KC depletion on PentV-induced hepatic capture of serotype-15A pneumococci was assessed as in Fig. 5B. Scale bar, 20 μm. n = 6-12. **(D** to **F)** Contribution of Fcγ receptors and the complement system to PentaV-induced capture of serotype 15A pneumococci wa5s assessed as in Fig. 5D-5G. (**G**) The types of antibodies against serotype-4 capsule or individual CPPs in mouse immune sera were detected by ELISA. n = 6. The representative data of at least two independent experiments are presented as mean ± SD, with statistical differences determined by unpaired *t* test (A), two-way ANOVA with Tukey’s multiple comparisons test (D), and multiple *t* tests (B, C, E and F). ns, no significant difference; \*\**P* < 0.01; \*\*\**P* < 0.001; \*\*\*\**P* < 0.0001.

**Fig. S6.**
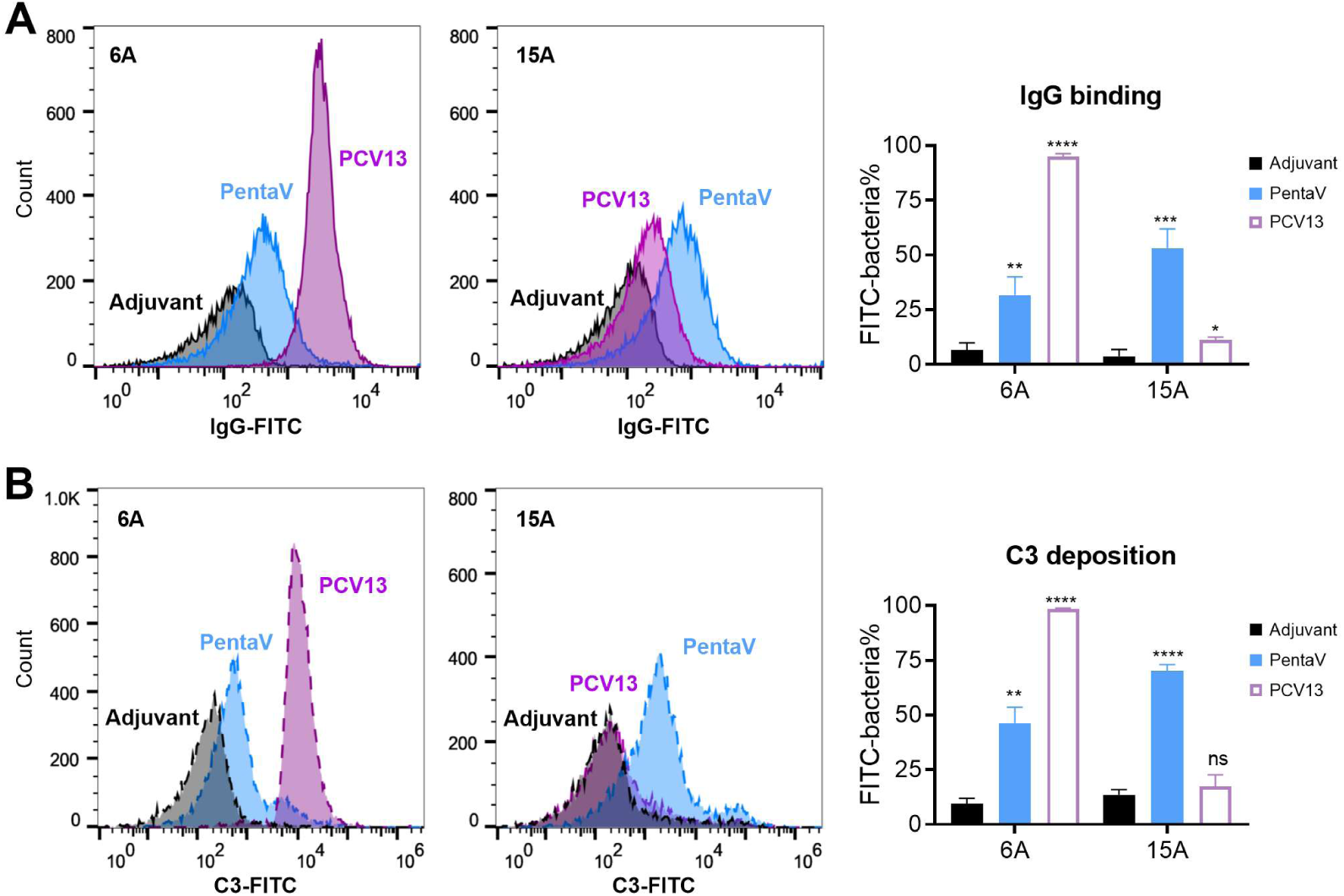
IgG and C3 deposition on serotype-6A and -15A pneumococci. IgG (A) and C3 (B) deposition on serotype-6A and -15A pneumococci were detected by flow cytometry after incubation with pentavalent protein vaccine or PCV13 immune serum as in Fig. 6. The percentages of IgG- or C3-positive bacteria are shown in the right panel. n = 3. The representative data of at least two independent experiments are presented as mean ± SD, with statistical differences determined by multiple *t* tests. ns, no significant difference; \**P* < 0.05; \*\**P* < 0.01; \*\*\**P* < 0.001; \*\*\*\**P* < 0.0001.

**Fig. S7.**
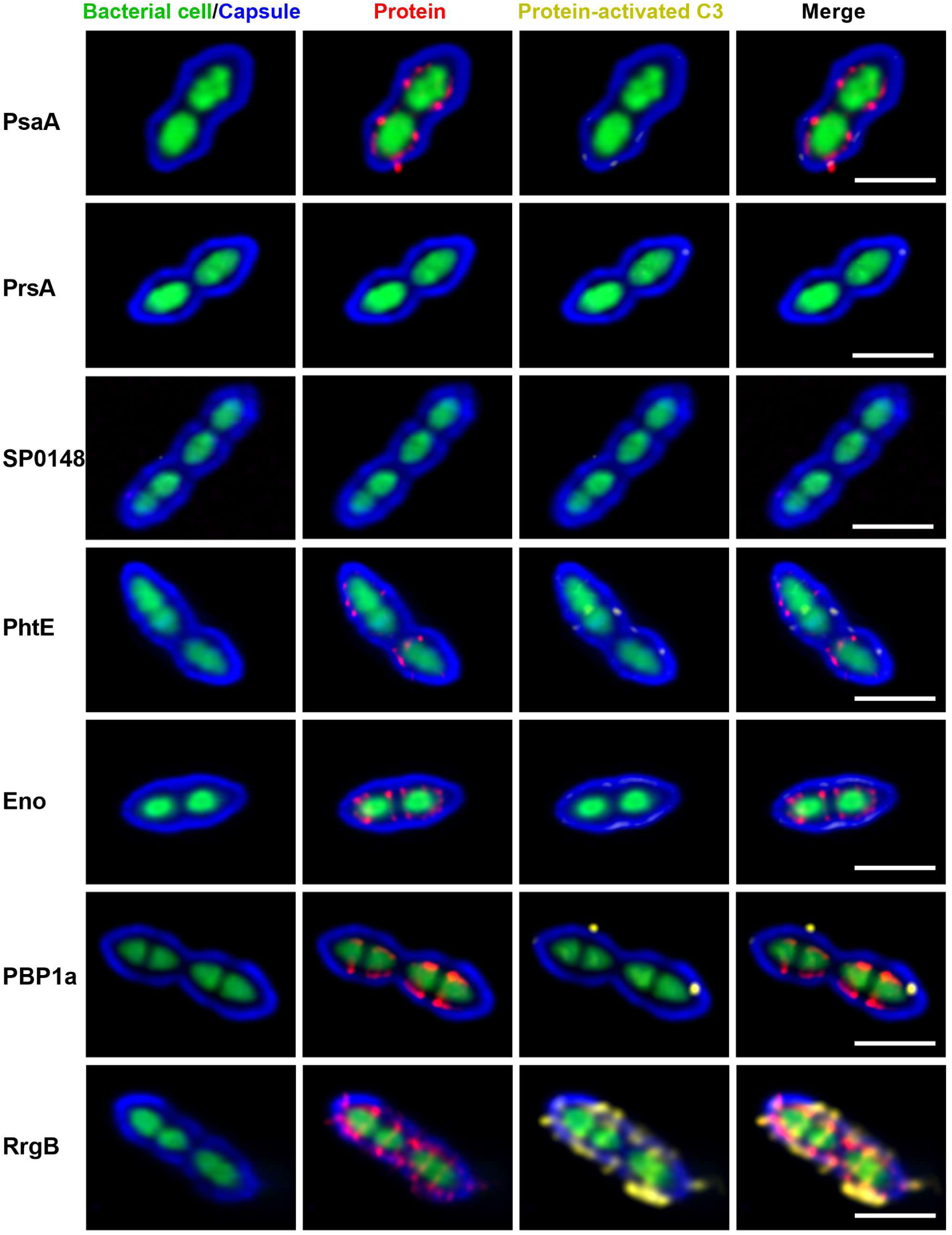
Spatial localization of protein-induced IgG and C3 on encapsulated bacteria. The relative positions of protein-specific IgG (red) and C3(yellow) on GFP-expressing pneumococci (green) were visualized after treating with immue sera as in Fig. 7. Scale bar, 2 μm. Representative data of at least two independent experiments are shown.

